# Structural elucidation of how ARF small GTPases induce membrane tubulation for vesicle fission

**DOI:** 10.1101/2023.12.19.572083

**Authors:** Xiaoyun Pang, Yan Zhang, Kunyou Park, Zhenyu Liao, Jian Li, Jiashu Xu, Minh-Triet Hong, Guoliang Yin, Tongming Zhang, Yaoyu Wang, Edward H. Egelman, Jun Fan, Seung-Yeol Park, Victor W. Hsu, Fei Sun

**Affiliations:** National Key Laboratory of Biomacromolecules, CAS Center for Excellence in Biomacromolecules, Institute of Biophysics, Chinese Academy of Sciences, Beijing 100101, China; Department of Life Sciences, Pohang University of Science and Technology, Pohang, Gyeongbuk 37673, Republic of Korea; Division of Rheumatology, Inflammation and Immunity, Brigham and Women’s Hospital, and Department of Medicine, Harvard Medical School, Boston, MA 02115 USA; School of Life Sciences, University of Chinese Academy of Sciences, Beijing 100049, China; Department of Biochemistry and Molecular Genetics, University of Virginia, Charlottesville, Virginia, VA 22908 USA; Center for Biological Imaging, Institute of Biophysics, Chinese Academy of Sciences, Beijing 100101, China; City University of Hong Kong, Hong Kong, China

## Abstract

The ADP-Ribosylation Factor (ARF) small GTPases have been found to act in vesicle fission through a direct ability to tubulate membrane. Here, we have used cryo-electron microscopy (EM) to solve the structure of an ARF6 protein lattice assembled on tubulated membrane to 3.9 Å resolution. ARF6 forms tetramers that polymerize into helical arrays to form this lattice. We identify, and confirm functionally, protein contacts critical for this lattice formation. The solved structure also suggests how the ARF amphipathic helix is positioned in the lattice for membrane insertion, and how a GTPase-activating protein (GAP) docks onto the lattice to catalyze ARF-GTP hydrolysis in completing membrane fission. As ARF1 and ARF6 are structurally conserved, we have also modeled ARF1 onto the ARF6 lattice, which has allowed us to pursue the reconstitution of Coat Protein I (COPI) vesicles to confirm more definitively that the ARF lattice acts in vesicle fission. Our findings are notable for having achieved the first detailed glimpse of how a small GTPase bends membrane and having provided a molecular understanding of how an ARF protein acts in vesicle fission.

## INTRODUCTION

The ADP-Ribosylation Factor (ARF) family of small GTPases acts in intracellular transport, a fundamental cellular process that is critical for the proper localization of proteins and lipids within the cell. Early studies identified a regulatory role for ARF proteins, which involves their recruitment of coat complexes onto membrane compartments to initiate the formation of vesicular carriers ^1,2^.

This role has been best characterized for ARF1 and an ARF-like protein known as Sar1. At the Golgi, ARF1 recruits the Coat Protein I (COPI) complex to form vesicular carriers for transport from the Golgi to the endoplasmic reticulum (ER) ^3,4^. ARF1 also recruits adaptor proteins (AP), which couple with clathrin in some cases, for transport from the Golgi to endosomal compartments ^5–7^. At the ER, Sar1 recruits the COPII complex for transport from the ER to the Golgi ^8^.

ARF6 has also been suggested to act in coat recruitment, in this case regulating a clathrin complex that uses ACAP1 (ArfGAP with Coiled-Coil, Ankyrin Repeat and PH Domain 1) and Akt as co-adaptors for transport from the recycling endosome to the plasma membrane ^9,10^. However, in contrast to the case of COPI regulated by ARF1 and COPII regulated by Sar1, for which vesicle formation has been reconstituted to enable detailed mechanistic studies ^4,8^, a similar achievement for the ARF6-regulated coat has not been attained.

A molecular understanding of how ARF proteins recruit coat complexes to membrane compartments is being achieved through structural studies. Co-crystal structures of ARF1 in complex with subunits of coatomer (the core components of the COPI complex) ^11^, and Sar1 in complex with Sec23/24 (the inner components of the COPII complex) ^12^, have revealed how these prototypic ARF proteins interact with their cognate coats for membrane recruitment.

Besides having a regulatory role, ARF proteins have also been found to play effector roles in vesicle formation. Vesicle formation occurs in two general stages, with vesicle budding involving positive membrane curvature and vesicle fission involving negative membrane curvature. Structural studies through high-resolution cryo-electron microscopy (EM) are providing a molecular understanding of how ARF1 assembles with coatomer on membrane for the budding stage of COPI vesicle formation ^13–15^. Similar studies are also providing molecular insights into how Sar1 assembles with the COPII components for bud formation ^16–18^.

Whereas these findings reveal how ARF proteins act together with their cognate coats to achieve membrane bending, a surprising finding has been that ARF proteins alone can also bend membrane. This ability was initially discovered by incubating Sar1 with liposomes generated from defined lipids, which resulted in liposome tubulation ^19^. Vesicle reconstitution studies then pinpointed this effect of Sar1 to be required for COPII vesicle fission ^19^. Similarly, ARF1 has been found to induce liposome tubulation, and the COPI vesicle reconstitution studies have revealed that this action of ARF1 is required for vesicle fission ^20^. However, in contrast to their roles in coat recruitment and vesicle budding, a structural understanding of how ARF proteins promote vesicle fission through membrane tubulation remains to be achieved.

In the current study, we address this notable gap in knowledge by initially solving the structure of an ARF6 protein lattice assembled on tubulated membrane to 3.9 Å resolution, which provides the first detailed glimpse of how a small GTPase bends membrane. Moreover, as the solved structure predicts five major protein interfaces that are critical for lattice formation, we have pursued functional mutagenesis studies to confirm these predictions. As the structures of ARF1 and ARF6 are highly conserved, we have also modeled ARF1 onto the ARF6 lattice, which has enabled us to pursue the COPI vesicle reconstitution system to confirm more directly that the ARF lattice structure acts in vesicle fission. A key approach involved mutating ARF1 residues needed to maintain the lattice structure and finding that COPI vesicle fission was inhibited. We also ruled out that these mutations affected instead the ability of ARF1 to interact with other proteins that act in COPI vesicle formation. Thus, we have achieved a molecular understanding of how an ARF protein acts in vesicle fission.

## RESULTS

As both ARF1 and ARF6 have been found previously to induce liposome tubulation ^21^, we performed pilot studies and found that the ARF6-induced tubules were more uniform. Thus, we pursued cryo-EM studies on these tubules.

### ARF6 lattice on tubular membrane

We first co-expressed ARF6 and N-myristoyl transferase in bacteria to generate myristoylated ARF6 (**Extended Data Fig. 1a and 1b**). We then confirmed that loading with GTP, but not GDP, enabled ARF6 to induce constricted tubules from liposomes with diameter of ∼ 30 nm (**Extended Data Fig. 1c**), similar to that previously observed ^21^.

We next vitrified samples that contain ARF6-induced tubules followed by imaging through cryo-EM. From the raw electron micrographs, we could readily observe the outer surface of tubulated liposomes to be coated with ARF6 proteins (**Extended Data Fig. 1d**). The power spectra of these tubules indicated that they belong to a 5-start helical assembly with a helical rise of 28.0 Å and twist of 27.3° (**Extended Data Fig. 1d**).

Combining the iterative helical real-space reconstruction approach^22^ with refinement using RELION ^23^ (**Extended Data Fig. 1d**), we determined the 3D density map of the ARF6 assembly on tubulated membrane to 3.9 Å resolution (**Extended Data Fig. 2a and 2b; Extended Data Table 1 and Supplementary Video 1**). We found that ARF6 forms a right-handed helical tubule with outer and inner diameters of 31.4 nm and 12.0 nm, respectively (**Fig. 1a and Extended Data Fig. 2c**).

**Fig. 1.**
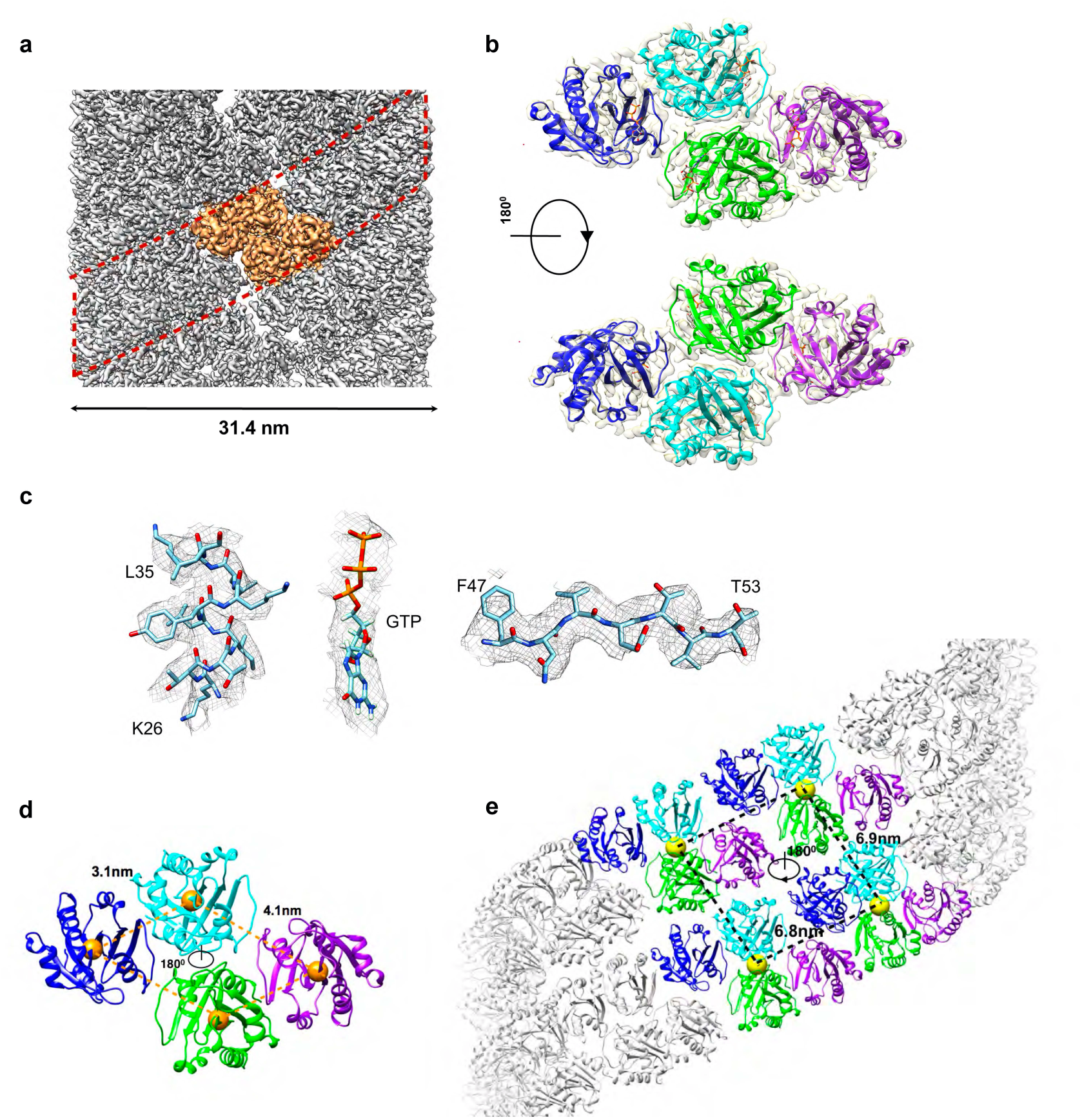
Cryo-EM reconstruction and atomic model of ARF6 coated tubule. **a**, Cryo-EM map of the reconstructed ARF6-coated tubule. A single helical rung (red dashed lines) and a helical asymmetric unit (orange) are highlighted. **b**, Model of ARF6 tetramer on membrane with each subunit shown in a different color: blue, cyan, green and purple. The bound GTP molecules are shown as sticks. **c**, Representative cryo-EM densities of side chains of α-helix 1 (K26-L35) (left), GTP molecule (center), and a strand from β-sheet 2 (F47-T53) (right). **d**, Tetramer organization. Centroid of each subunit is represented by an orange ball. Dotted lines that connect four adjacent centroids form a parallelogram. **e**, ARF6 tetramers in helical arrays. Centroid of each tetramer is represented by yellow ball. Dotted lines that connect four adjacent centroids form a rhombus.

We next sought to fit previously solved crystal structures of ARF6 onto the density map. Initial rigid body fitting by Chimera^24^ showed the cross correlation between the crystal structure and the experimental density to be 0.63 for ARF6 in its GTPγS-bound state [Protein Data Bank (PDB) ID: 2J5X] ^25^, and 0.48 in its GDP-bound state (PDB ID: 1E0S) ^26^. Thus, as ARF6-GTP provided the better fit, its atomic coordinates were then refined to fit on to the density map more precisely (**Fig. 1a and 1b**). This enabled many bulky side chains to be resolved (**Fig. 1c**). ARF6 binding to GTP in forming the lattice structure was further verified by the detection of densities that correspond to GTP binding (**Fig. 1c**).

The solved structure further revealed that four ARF6 molecules assemble into a tetramer with a two-fold symmetry. When the centroids of these four ARF6 molecules within the tetramer are connected, they form a parallelogram configuration (**Fig. 1d and Supplementary Video 2).** The lengths of the long sides and short sides of the parallelogram are 4.1 nm and 3.1 nm, respectively. This parallelogram-shaped tetramer is the asymmetric unit of helical packing (**Fig. 1a and 1e**). Each tetramer is oriented with the N-terminus of ARF6 facing toward the membrane and the GTP-binding pocket facing the opposite direction (**Supplementary Video 2**). Whereas connecting the centroids inside the tetramer form a parallelogram, connecting the centroids of four tetramers grouped from two adjacent helical rows form a rhombus. These tetramers interlock with each other to reinforce the overall architecture of the lattice (**Fig. 1e**).

### Protein interfaces critical for the ARF6 lattice

The solved structure predicted multiple protein interfaces critical for the ARF6 lattice assembly (**Fig. 2a and 2b, and Extended Data Table 2**). These interfaces can be divided into three types: i) interactions in forming the tetramer (interfaces 1, 2 and 3), ii) interactions between tetramers in the same helical row (interface 4), and iii) interactions between tetramers in adjacent helical rows (interface 5) that crosslink the helical rows (**Fig. 2a and 2b**).

**Fig. 2.**
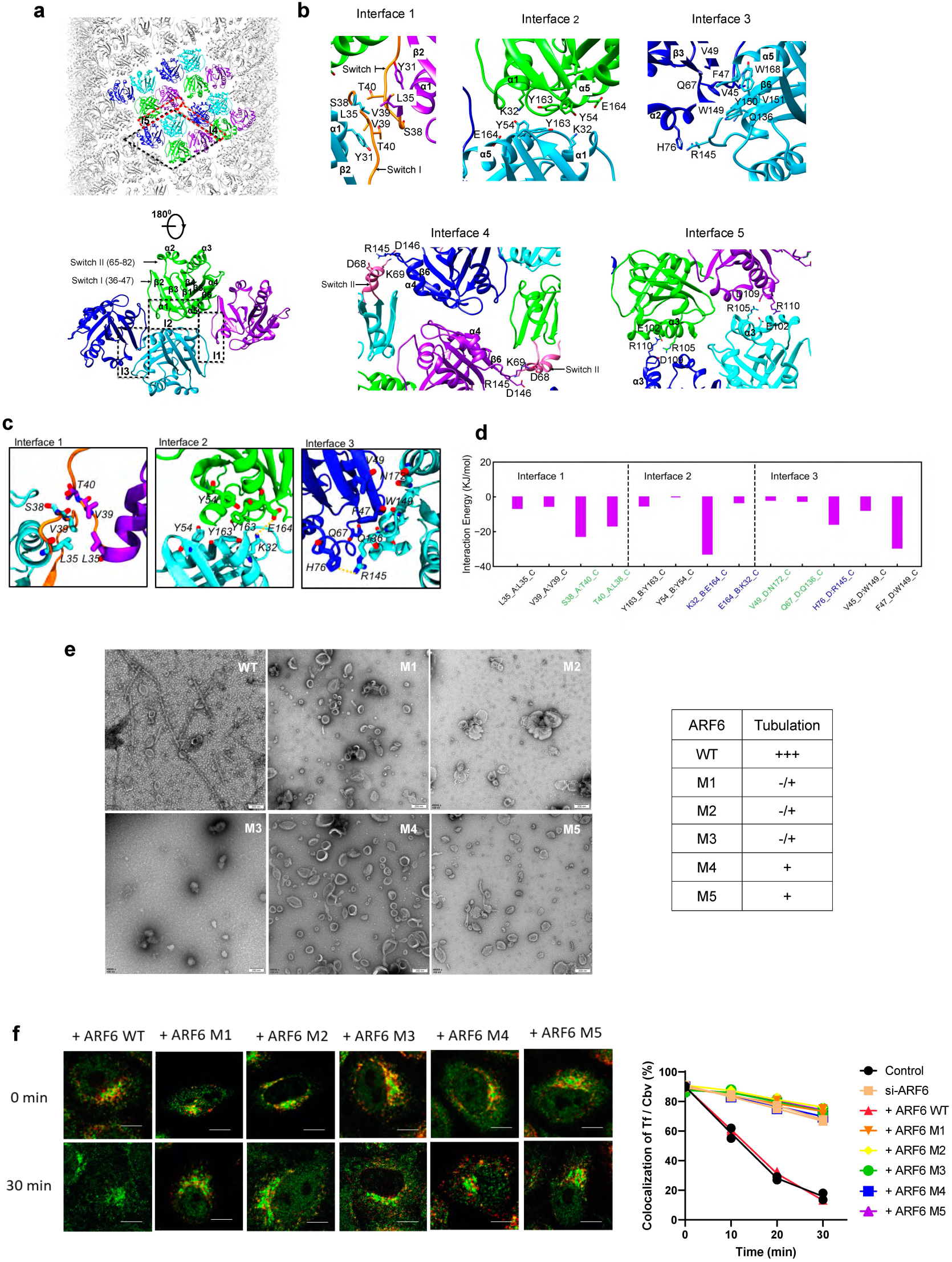
Protein interactions critical for the ARF6 lattice assembly. **a**, Organization of ARF6 tetramers. Upper panel: black dashed box outlines four subunits forming a tetramer, and red dashed box outlines two adjacent tetramers. Lower panel: close-up of the tetramer assembly with secondary structures and the major protein interfaces shown. **b,** Close-up of the major protein interfaces in the tetramer. Interface 1 is formed by helix α1 and switch I interacting with their counterpart in an adjacent ARF6 molecule in an anti-parallel manner. Switch I is highlighted in orange. Interface 2 is formed by helix α5 and inter-switch from adjacent molecules forming a symmetric interaction. Interface 3 is formed by switch I, sheet β6 and helix α5 between two adjacent molecules forming interactions. Interface 4 involves tetramer interactions within the same helical row, highlighting the role of two pairs of electrostatic interactions (D68-R145 and K69-D146). Switch II is highlighted in pink. Interface 5 involves tetramers on different helical rows interacting with each other, highlighting the role of salt bridges (E102-R110, R105-D109) forming symmetric interactions. **c**, Snapshots of the major protein interfaces of the membrane-bound ARF6 tetramer at the final frame of the molecular dynamics simulations. Salt bridges are indicated by yellow dashed lines. **d**, Interaction energy for each residue pair in the protein interfaces, with salt bridges and hydrogen bonds highlighted by green and blue labels, respectively. **e**, EM analysis examining the effect of ARF6 mutations on liposome tubulation. Specific residues mutated are: M1, Y31A/L35A/V39A; M2, Y54A/Y163A; M3, V45A/F47A/W149A/Y150A/V151A /W168A; M4, K69E/R145E; M5, R105E/R110E. Representative images are shown on left; scale bar, 200 nm. Quantitation of tubulation ability is shown on right: +++ (strong), ++ (moderate), + (weak), or - /+ (almost none). **f,** The TfR recycling assay assessing the effect of ARF6 mutations on transport from the recycling endosome to the plasma membrane (PM). Representative images of endosomal TfR (red) colocalizing with a recycling endosome marker, cellubrevin (Cbv, green), at times indicated are shown on left. Quantitation is shown on right, comparing control, siRNA against ARF6, and rescues using different ARF6 mutants.

Interface 1 occurs through an anti-parallel interaction between two ARF6 molecules in forming a dimer, and involves helix α1 and its connecting loop, namely Switch I (residues 36–47) between helix α1 and sheet β2 of one subunit interacting with that of another subunit. Interactions in this interface are mediated by hydrogen bonds (S38-T40) and hydrophobic contacts (Y31, L35 and V39) (**Fig. 2b**).

Interface 2 is formed by interaction between two opposing ARF6 molecules that have a two-fold symmetry and involves the tip of Interswitch I (residues 48-64) and helix α5. Hydrophobic residues Y163 and Y54 from these two subunits form a symmetric interaction (**Fig. 2b**). Additional salt bridges (K32-E164) between helix α1 and helix α5 from these neighboring subunits further stabilize the interface (**Fig. 2b**).

Interface 3 is formed by two ARF6 interacting through Switch I, β6 sheet, and α5 helix. Hydrophobic residues V45 and F47 of one subunit interact with residues W149, Y150, V151 and W168 of an adjacent subunit. Additional salt bridge (H76-R145) and hydrogen bonds (Q67-Q136 and V49-N172) from these neighboring ARF6 molecules further stabilize the interface (**Fig. 2b**).

Interface 4 is formed by two adjacent ARF6 tetramers along the same helical row interacting with each other through short contacts. This involves the loop in Switch II (residues 65–82, between sheet β3 and helix α2) and the loop between helix α4 and sheet β6 of one subunit interacting with those of another subunit. This interface is likely supported by electrostatic interactions (D68-R145 and K69-D146) (**Fig. 2b**).

Interface 5 is formed by ARF6 tetramers in adjacent helical rows interacting with each other with a two-fold symmetry. This involves α3 and η3 helices of a tetramer residing in one helical row interacting with another in a tetramer residing in an adjacent row (**Fig. 2b**). Salt bridges (E102-R110, R105-D109) form symmetric interactions between tetramers that reside in adjacent helical rows.

As the tetramer is the basic unit of the ARF1 lattice structure, we next pursued molecular simulation studies to further characterize the three major interfaces predicted to be critical for its assembly (**Fig. 2c and 2d**). Simulation was initiated by placing the ARF6 tetramer on top of the membrane (**Supplementary Video 3**). For Interface 1, the hydrophobic contacts (L35 and V39) and the hydrogen bond (S38-T40) remain stable throughout the simulation. The interaction energies of hydrogen bonds between two pairs of S38 and T40 were similar, consistent with Interface 1 having anti-parallel interactions. For Interface 2, simulation highlighted particularly the importance of the salt bridge (K32-E164), as it had the highest calculated interaction energy among the predicted interactions in this interface. For Interface 3, simulation confirmed the importance of salt bridges (H76-R145) and particularly the hydrophobic interaction between F47 and W149, as they had the highest calculated interaction energies.

### Functional mutagenesis studies targeting the protein interfaces

We then pursued functional mutagenesis studies to further confirm the importance of the predicted major interfaces. For interface 1, we targeted residues Y31, L35, and V39 (**Fig. 2b**). For interface 2, we targeted residues Y163 and Y54 (**Fig. 2b**). For interface 3, we targeted residues V45, F47, W149, Y150, V151 and W168 (**Fig. 2b**). These residues were all mutated to alanines to eliminate hydrophobic interactions. The resulting mutants were named M1, M2, and M3, respectively. For interfaces 4 and 5, predicted key residues are all charged residues. Thus, we mutated positively charged residues (lysine and arginine) to glutamates. To target interface 4, we mutated K69 and R145 (**Fig. 2b**) and the resulting mutant was named M4. For interface 5, we mutated R105 and R110 (**Fig. 2b**) and the resulting mutant was named M5.

We initially performed circular dichroism (CD) spectroscopy, which revealed that the mutants maintained similar conformations as that of the wild-type form (**Extended Data Fig. 2d**). We then performed liposome studies to confirm that targeting against the interfaces reduced the ability of ARF6 to induce liposome tubulation (**Fig. 2e**). As ARF6 acts in endocytic recycling, we also confirmed that this transport was inhibited by the mutations, which was tracked by the recycling of the transferrin receptor (TfR) from the recycling endosome to the plasma membrane (**Fig. 2f**), as previously described ^10^.

### ARF6 lattice is distinct from those formed by other proteins

We next compared the ARF6 lattice to those previously solved for other membrane-bending proteins. These included proteins that contain the BAR domain, such as endophilin ^27^, ACAP1 ^28^, and SNX1 ^29^, as well as dynamin that does not contain a BAR domain ^30^. The result revealed distinct features in the ARF6 lattice structure. ARF6 assembles as a 5-start helical array. In contrast, the BAR domain of endophilin (N-BAR), as well as dynamin in its GTP-bound state, are 2-start helical assemblies, while SNX1 and the BAR-PH tandem domain of ACAP1 (BAR-PH) assemble as a 1-start helical array (**Extended Data Fig. 2e**).

Another distinction was that the asymmetric unit of the ARF6 assembly is a tetramer, while those of endophilin, SNX1 and dynamin are dimers, and that of ACAP1 is a dimer of dimer (**Extended Data Fig. 2e**). These differences result in the ARF6 lattice having an overall architecture that could be readily distinguished from lattices formed by the other proteins, as seen in the cross-section views of the different coated tubules (**Extended Data Fig. 2f**). Thus, the ARF6 lattice represents a new type of protein interaction and assembly pattern that mediates membrane bending.

### Positioning of the amphipathic helix in the ARF6 lattice

We next considered that ARF proteins have been found to contact the membrane through two structural elements, an N-terminal myristoyl chain and an amphipathic helix located also near the N-terminus ^1,2,21,31^. In the solved lattice structure, the myristoyl chain merges imperceptibly with the lipids of the membrane. However, the structure of the amphipathic helix could be readily discerned.

We performed a low-pass filter of the cryo-EM map to 7 Å. At this resolution, additional densities connecting N11 were displayed clearly which arise from the N-terminal helix (**Fig. 3a**). Docking the helix onto the ARF6 lattice suggested that it inserts substantially into the membrane (**Fig. 3b**). As the helix had been predicted to be amphipathic ^21^ with the hydrophobic face consisting of V4, L5, I8 and F9, and the hydrophilic face consisting of K3, S6, and K7, our confirmation of this orientation of the helix within the membrane further supported the validity of how the lattice structure is assembled on membrane (**Fig. 3b**).

**Fig. 3.**
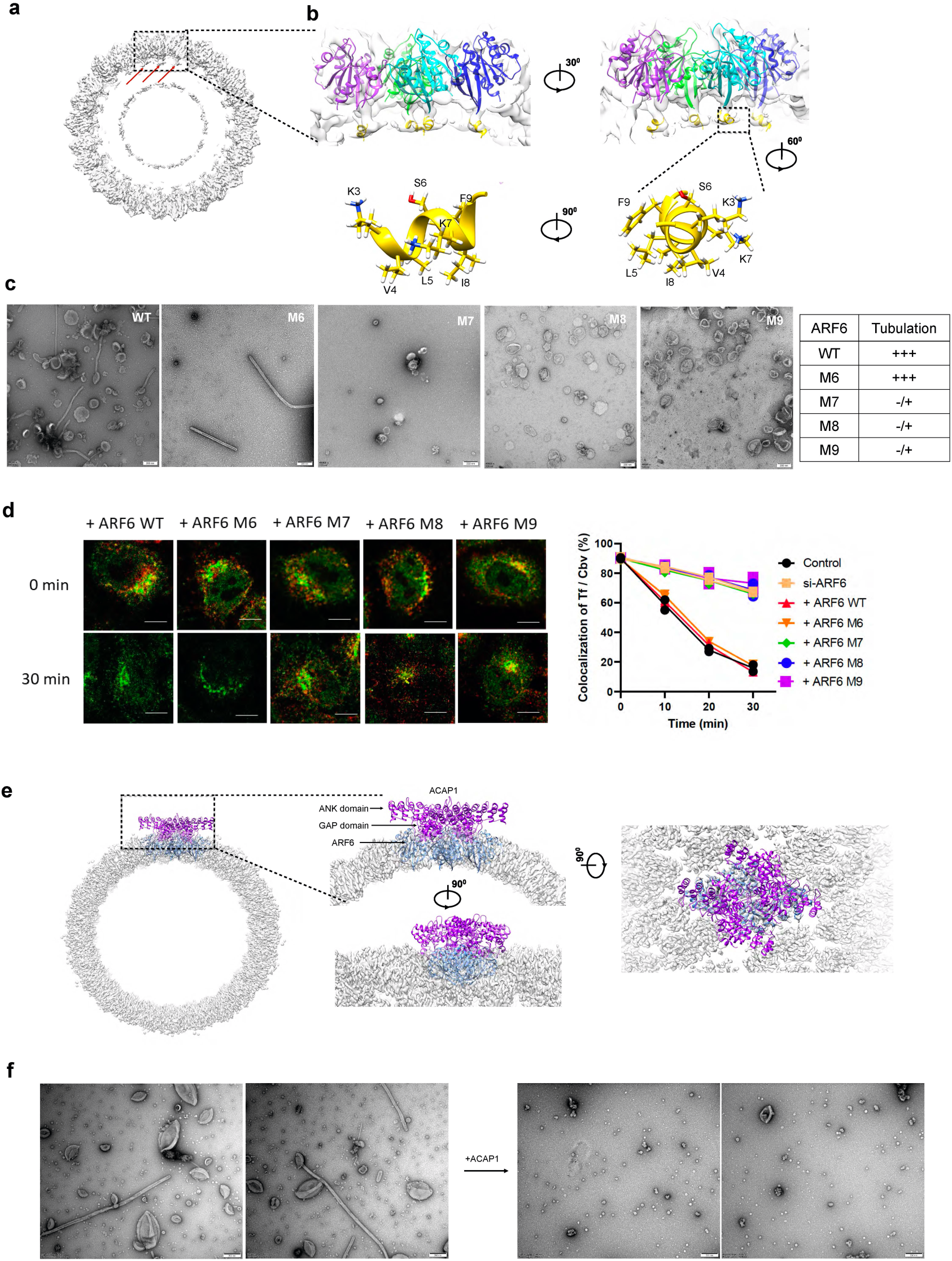
Interaction of the ARF6 lattice with membrane tubule. **a**, Cross-sectional view of reconstructed ARF6-coated tubules. Black dashed box outlines a tetramer cluster. **b**, Close-up views of the ARF6 lattice interacting with the membrane. Upper panels show a tetramer with its subunits (in four colors) interacting with the membrane, either in the same orientation as the cross-sectional view or rotated by 30°. Lower panels show the ribbon model of the amphipathic helix with charged face on one side and hydrophobic face on the opposite side. Key residues on both faces are labeled. **c,** EM analysis examining the effect of ARF6 mutations on liposome tubulation. Specific residues mutated are: M6, F9W; M7, F9A; M8, F9E; M9, V4E/L5E/I8E. Representative images are shown; scale bar, 200 nm. Quantitation of tubulation ability is shown as a table: +++ (strong), ++ (moderate), + (weak), or - /+ (almost none). **d,** The TfR recycling assay assessing the effect of ARF6 mutations on transport from the recycling endosome to the plasma membrane. Representative images of endosomal TfR (red) colocalizing with Cbv (green), at times indicated are shown on left. Quantitation is shown on right, comparing control, siRNA against ARF6, and rescues using different ARF6 mutants. **e**, The GAP and ANK domains of ACAP1 can be docked onto the ARF6 lattice structure without structural collision. **f,** Incubation of ACAP1 (GAP and ANK domains) with ARF6 liposomal tubules results in vesiculation.

To further characterize how the tetramer engages the membrane, we next pursued molecular dynamics simulation studies (**Supplementary Video 3**). The results confirmed that ARF6 tetramer associates firmly with the lipid membrane through the N-terminal myristoyl chain (MYR) and amphipathic helix (AH) (**Extended Data Fig. 3a**), with the latter involving the insertion of hydrophobic residues V4, L5, I8 and F9 into the membrane. The simulation also provided a temporal description of how the MYR, AH, as well as other parts of the tetramer engage the membrane (**Extended Data Fig. 3b**). The time evolved separation profile of molecular dynamics confirmed that the MYR and AH regions fully insert into the membrane (**Extended Data Fig. 3c**). Moreover, the hydrophobic face of AH penetrates the membrane more deeply than the hydrophilic face (**Extended Data Fig. 3a**). The insertion of the region is further supported by its reduction in root mean square fluctuation (RMSF) upon membrane binding (**Extended Data Fig. 3d**).

We next pursued functional mutagenesis to validate our structural predictions. Focusing on the aromatic residue F9 in the amphipathic helix, we found that a conservative mutation to tryptophan (referred as M6) largely retained liposome tubulation by ARF6 (**Fig. 3c**). In contrast, mutation to alanine (referred as M7) or to glutamate (referred as M8) inhibited tubulation. When multiple residues that line the hydrophobic face of the amphipathic helix were mutated to glutamates (V4E, L5E, and I8E; referred as M9), tubulation was also reduced (**Fig. 3c**). Consistent with these findings, we found that M6 did not have a significant effect on TfR recycling, while the other mutations (M7, M8, M9) inhibited this recycling (**Fig. 3d**).

### Interaction of ARF GAPs with the ARF6 lattice

The GAPs that act on ARF proteins have also been found to act in vesicle fission for the better characterized coats ^32,33^. Thus, we next examined whether the solved ARF6 lattice structure would allow the GAP to dock onto the lattice to catalyze GTP hydrolysis on ARF6. An ARF6-GAP complex has been solved previously, which involves ARF6 bound to ASAP3 (PDB ID: 3LVQ) ^34^. The overall structure of ARF6, when complexed with ASAP3, is similar to that of our ARF6 assembled on membrane, with the r.m.s.d between the two being 0.6 Å for 164 C_α_ atoms. After superimposing the structure of the ARF6-ASAP3 complex onto the ARF6 lattice structure, ASAP3 is predicted to be located on the surface of lattice assembly with its GAP domain positioned toward the active site of ARF6, with no structural collisions predicted for this configuration (**Extended Data Fig. 3e**). Thus, the ARF6 lattice structure should allow the subsequent action of the GAP activity to promote vesicle fission.

Further supporting this prediction, we performed a similar docking for ACAP1, the GAP that acts on ARF6 in endocytic recycling ^9^, and found that its GAP domain would also be positioned on the ARF6 lattice structure to allow ARF-GTP hydrolysis (**Fig. 3e**). As functional confirmation, we then incubated ACAP1 with the ARF6-induced liposomal tubules, and notably, vesiculation was observed (**Fig. 3f**). Thus, the results not only confirmed that the lattice structure would allow the GAP to act for ARF-GTP hydrolysis, but also predicted that vesicle fission would involve the sequential actions of ARF-induced constriction of the bud neck followed by GAP-induced neck scission.

### Modeling ARF1 onto the ARF6 lattice

We then sought to confirm more directly that the ARF lattice structure acts in vesicle fission by pursuing vesicle reconstitution studies. A practical consideration was that this approach has not been estabished for the ARF6-regulated coat. Instead, because it has been established for vesicle formation by the COPI complex ^35,36^, we next examined whether ARF1 could be modeled onto the ARF6 lattice structure.

The feasibility of this modeling was suggested by the high degree of sequence homology between ARF1 and ARF6 (**Extended Data Fig. 4a**), and the r.m.s.d between the crystal structure of ARF1 in its GTP-bound state ^37^ (PDB ID 1O3Y) and the cryo-EM structure of ARF6 in its GTP-bound state (this study), which revealed that it is 1.6 Å for 163 C_α_ atom and 0.7 Å for 157 pruned C_α_ atom (**Extended Data Fig. 4b**). Further supporting the robustness of this modeling, we found that it predicted that the tetramer would also be the asymmetric unit of the ARF1 lattice on membrane tubules (**Extended Data Fig. 4c**). Moreover, conservation in structure extended to each of the predicted protein interfaces (**Extended Data Fig. 4d**).

For ARF1, interface 1 was predicted to be formed by the anti-parallel (close to C2 symmetry) interaction of hydrogen bonds (I42-T44) and hydrophobic contacts (Y35, L39 and V43) (**Extended Data Fig. 4d**). Interface 2 would require conserved hydrophobic residues Y167 and Y58 from adjacent subunits interacting with each other, with salt bridges (K36-E168) further stabilizing this interface (**Extended Data Fig. 4d**). For interface 3, hydrophobic residues I49, F51 and V53 of one subunit, were predicted to interact with W153, Y154, I155 and W172 of an adjacent subunit (**Extended Data Fig. 4d**). The same interactions between the other paired subunits would further contribute to stabilize the tetramer organization. For interface 4, two adjacent ARF1 tetramers would interact with each other along the same helical row through short contacts, likely formed by electrostatic interactions (D72-R149 and K73-H150, **Extended Data Fig. 4d**). Interface 5 was predicted to involve contacts between adjacent helical rows, in which salt bridges (R109-E113) would form symmetric interactions between tetramers on different helical rows (**Extended Data Fig. 4d**).

### Elucidating the role of the ARF1 lattice in COPI vesicle formation

To confirm these predictions, we again pursued functional mutagenesis. We generated mutations in ARF1 that paralleled those done for ARF6. For Interface 1, we mutated residues I42 and T44 in ARF1 to alanines. For interface 2, we mutated residues Y167 to alanine and E168 to arginine. For interface 3, we mutated residues I49 and F51 to alanines. The resulting mutants were named ARF1-M1, ARF1-M2, and ARF1-M3, respectively. For interface 4, we mutated K73 and R149 to glutamates and this mutant was named ARF1-M4. For interface 5, we mutated R109 to glutamate and D114 to arginine and this mutant was named ARF1-M5 (**Extended Data Fig. 4b**). Moreover, to target the amphipathic helix, we mutated the F13 residue (corresponding to F9 of ARF6, see **Extended Data Fig. 4a**) on the hydrophobic face to glutamate (F13E), and this mutant was named ARF1-M6.

Initially, to assess the physiologic relevance of these mutations, we performed the COPI transport assay in cells, which involved tracking a model COPI cargo, known as VSVG-KDELR, in its redistribution from the Golgli to the ER, as previously described ^38^. The results confirmed that the ARF1 mutations all inhibited COPI transport (**Fig. 4a and 4b**).

**Fig. 4.**
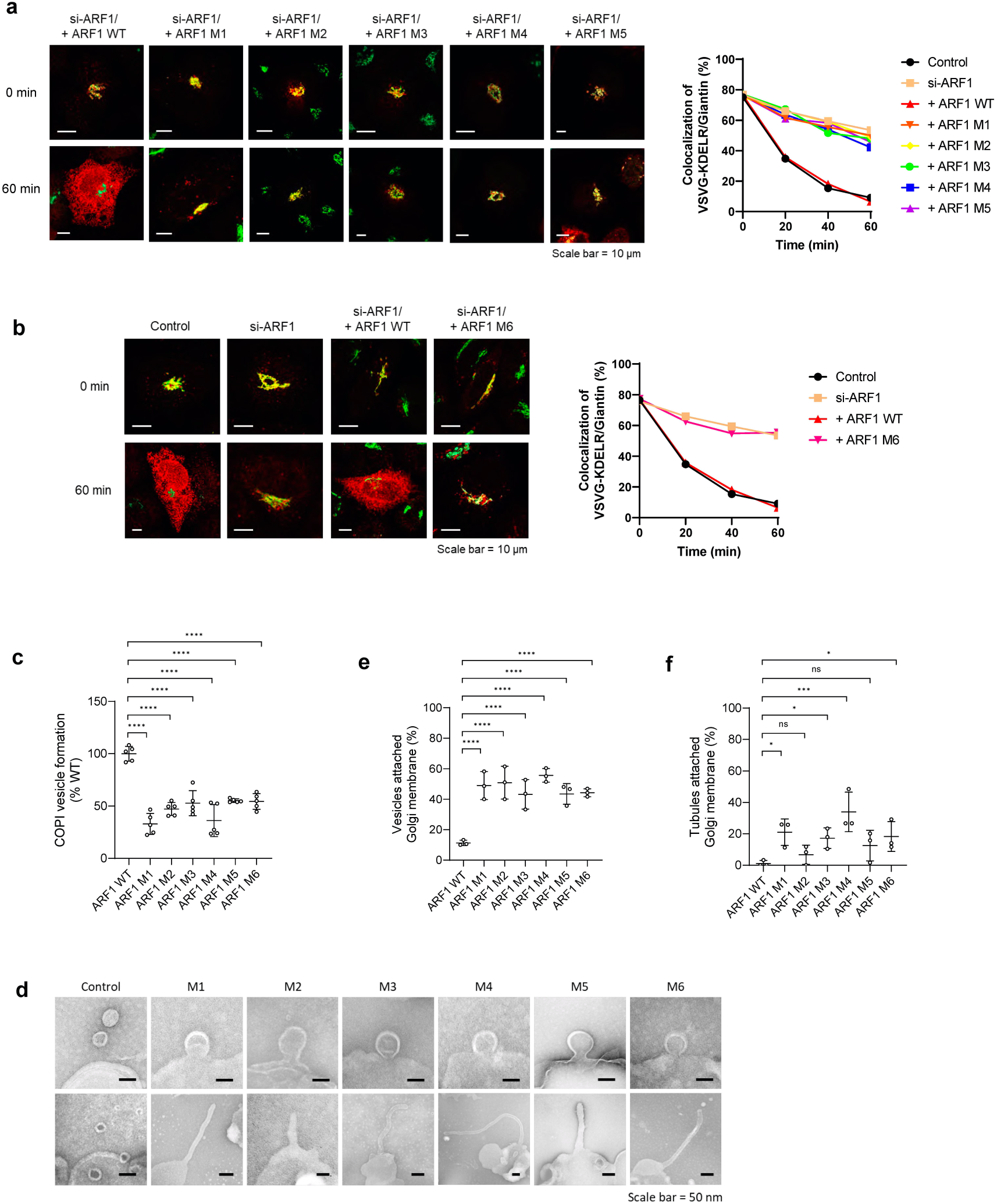
The role of the ARF1 lattice structure in COPI transport. **a**, COPI transport assay examining the effect of ARF1 mutations. Specific residues mutated are: M1, I42A/T44A; M2, Y167A/E168R; M3, I49A/F51A; M4, K73E/R149E; M5, R109E/D114R. Representative images of VSVG-KDELR (red) colocalizing with a Golgi marker (giantin, green) at times indicated are shown on left. Quantitation is shown on right, comparing control, siRNA against ARF1, and rescues using different ARF1 forms as indicated. **b**, COPI transport assay examining the effect of a mutation that targets the amphipathic helix of ARF1 (M6, F13E). Representative images of VSVG-KDELR (red) colocalizing with giantin (green) at times indicated are shown on left. Quantitation is shown on right, comparing control, siRNA against ARF1, and rescues using different ARF1 forms as indicated. **c**, Reconstitution of COPI vesicles from Golgi membrane examining the effect of ARF1 mutations on vesicle formation. Quantitation from five independent experiments is shown. Values are expressed as mean with standard deviation. Statistical analysis was performed using the Student’s t-test, comparing the wild-type and different mutants, ****P < 0.0001. **d**, Representative EM images from the COPI vesicle reconstitution system showing arrest in the two stages of vesicle fission induced by the ARF1 mutations. Upper panels show arrest in late fission, as reflected by buds with markedly constricted necks. Lower panels show arrest in early fission, as reflected by the formation of tubules; scale bar, 50nm. **e**, Quantitation of COPI buds with constricted necks induced by using different ARF1 mutants in the vesicle reconstitution system. The mean with standard deviation from three independent experiments is shown. Statistical analysis was performed using the Student’s t-test, comparing the wild-type and different mutants, ****P < 0.0001. **f**, Quantitation of COPI tubules induced by using different ARF1 mutants in the vesicle reconstitution system. The mean with standard deviation from three independent experiments is shown. Statistical analysis was performed using the Student’s t-test, comparing the wild-type and different mutants, ****P < 0.0001.

We next pursued the reconstitution of COPI vesicle formation using Golgi membrane, as previously described ^33^. In recent years, this reconstitution has been refined to enable COPI vesicle fission to be subdivided into two stages, with the inhibition of early fission that prevents the constriction of the COPI bud neck resulting in the generation of COPI tubules, and the inhibition of late fission resulting in the accumulation of COPI buds with constricted necks ^39^. We initially confirmed that the ARF1 mutations which target each of the protein interfaces in the lattice structure, as well as the amphipathic helix, inhibited COPI vesicle formation (**Fig 4c**). We then performed EM examination and found that targeting the protein interfaces or the amphipathic helix resulted in mixed accumulations of tubules and buds with constricted neck (**Fig 4d**), predicting that the ARF1 lattice would participate in both the early and the late stages of COPI vesicle fission.

We then performed quantitation and found that the mutations had relatively uniform effect in accumulating COPI buds with constricted necks (**Fig 4e**), suggesting that the different protein interfaces and the amphipathic helix all contribute similarly to late fission. In contrast, the effect of the mutations in accumulating COPI tubules was more varied, with M2 having less effect and M4 having more effect (**Fig 4f**), predicting that the protein interface targeted by M2 would be less critical for early fission, while the interface targeted by M4 would be more critical for early fission.

### Examining whether the mutations affect ARF1 interacting with other proteins that act in COPI vesicle formation

A potential caveat of our conclusions above was that the mutations could be disrupting instead the ability of ARF1 to interact with other proteins needed for COPI vesicle formation. To address this possibility, we noted that how ARF1 interacts with coatomer on COPI vesicles has been elucidated in detail through cryo-EM studies ^14,15,40^. Thus, we superimposed our model of Arf1 tetramer onto the cryoEM structure of the COPI coat on these vesicles ^15^ (PDB ID, 5NZR). When an ARF1 subunit in the tetramer was aligned with an ARF1 subunit in the ARF1-coatomer complex, severe structural collision was observed between another ARF1 subunit in the tetramer and γ-COP within the ARF1-coatomer complex (**Extended Data Fig. 4e**).

We then systematically investigated all the interaction interfaces between ARF1 and coatomer subunits, which are β-COP, δ-COP, β’-COP, and γ-COP, in the ARF1-coatomer complex of the COPI coat and compared them with all the interaction interfaces within the ARF1 lattice structure.

We had mutated residues I42 and T44 to target Interface 1 of the ARF1 lattice. Although these residues were within the vicinity of the interfaces between ARF1 and δ-COP (**Extended Data Fig. 5b)** and between ARF1 and β’-COP (**Extended Data Fig. 5e)** in the ARF1-coatomer complex, the targeted ARF1 residues were not predicted to be directly involved in the two ARF1-coatomer interfaces. Instead, the interface between ARF1 and δ-COP was predicted to involve mainly E164 of ARF1 with E165 of δ-COP (potential hydrogen bond or salt bridge), L37 of ARF1 with E158/K162 of δ-COP (potential hydrogen bond with main chain of L37), and E54 of ARF1 with Q151 of δ-COP (potential hydrogen bond) (**Extended Data Fig. 5b)**, while the interface between ARF1 and β’-COP was predicted to involve mainly T161 of ARF1 with G173 of β’-COP, E41 of ARF1 with K219 of β’-COP, and E41 of ARF1 with N178 of β’-COP (**Extended Data Fig. 5e).**

To target Interface 2 of the ARF1 lattice, we had mutated residues Y167 and E168 to alanine and arginine, respectively. These residues were close to but also unlikely to participate in the interface between ARF1 and δ-COP in the ARF1-coatomer complex (**Extended Data Fig. 5b)**, as this interface was predicted to involve instead the paired residues noted above.

To target Interface 3 of the ARF1 lattice, we had mutated residues I49 and F51 to alanines. We noted the interface between Arf1 and γ-COP is predicted to involve mainly residues F51, L77, H80 and Y81 of ARF1 and residues I103, I104 and K57 of γ-COP, respectively (**Extended Data Fig. 5d and 5f),** while the interface between Arf1 and β-COP likely involves residues I46, F51, R73 and H80 of ARF1 interacting with residues L82, R79, M75, I114 and M115 of β-COP, respectively (**Extended Data Fig. 5a)**. Therefore, because there are multiple residues besides F51 that are involved in these interfaces, mutating only F51 seemed unlikely to affect the interactions between Arf1 and γ-COP/β-COP significantly.

To target Interface 4 of the ARF1 lattice, we had mutated residues K73 and R149 to glutamates. Whereas R149 was not within the vicinity of any of the ARF1-coatomer interfaces, K73 was situated in closer proximity to the interfaces between ARF1 and β-COP (**Extended Data Fig. 5a**, shown as R73 because the yeast ARF1 was used in the previous cryo-EM structure of ARF1-coatomer interactions) and between ARF1 and γ-COP (**Extended Data Fig. 5f**, shown as R73 for reason mentioned above) in the ARF1-coatomer complex. However, because K73 was situated at the edge of these interfaces, mutating this residue seemed unlikely to affect these ARF1-coatomer interfaces significantly.

To target Interface 5, we had mutated residues R109 and D114 to glutamate and arginine, respectively. These residues were more easily ruled out, as they are not even within the vicinity of any of the ARF1-coatomer interfaces. There is another interaction interface between Arf1 and γ-COP in the structural model of the ARF1-coatomer complex (**Extended Data Fig. 5c**), but this interface is also not within the vicinity of any ARF1 residues critical for the lattice formation.

Thus, as the results altogether predicted that ARF1 would be positioned quite differently in the lattice structure as compared to that in the ARF1-coatomer complex on COPI vesicles, mutations that target the lattice structure were unlikely to affect how ARF1 interacts with coatomer on COPI vesicles. Notably, this conclusion is further supported by a key observation. Whereas the ARF1-coatomer complex is predicted to play a central role in the budding stage of COPI vesicle formation, we had observed that the ARF1 mutations still allowed COPI vesicle formation to proceed through this stage, as only the fission stage was affected (see **Fig 4d-f**). Nevertheless, we also performed additional functional studies to further confirm that the ARF1 mutations would not affect its interaction with coatomer, as well as other proteins that act in COPI vesicle formation.

As the membrane recruitment of coatomer requires its interaction with ARF1, we found that the ARF1 mutations did not affect the ability of ARF1 to recruit coatomer onto the Golgi membrane (**Extended Data Fig. 6a)**. The mutations also did not affect the ability of ARF1 to be recruited to Golgi membrane (**Extended Data Fig. 6b)**, or the localization of ARF1 to the Golgi in cells (**Extended Data Fig. 6c)**. Thus, ARF1 interacting with factors needed for its recruitment to the Golgi was not affected by the mutations.

We further considered that, besides ARF1 and coatomer which are sufficient for vesicle formation from liposomes, COPI vesicle formation from Golgi membrane is more complex, requiring ARFGAP1 and BARS additionally ^36,41^. Interrogating the interaction between ARF1 and ARFGAP1, we found that it also was not affected by the ARF1 mutations (**Extended Data Fig. 6d)**. Furthermore, we found that ARF1 does not interact with BARS (**Extended Data Fig. 6e)**, and thus ruling out that this interaction could be a relevant target of the mutations. As such, these additional functional results further confirmed that the ARF1 mutations target the ARF1 lattice structure in explaining how they inhibit COPI vesicle fission.

## DISCUSSION

We have elucidated in molecular detail an ARF6 lattice structure assembled on tubulated membrane. To our knowledge, this structure represents the first detailed glimpse of how a small GTPase acts in membrane bending by forming a tubular protein lattice. The ARF6 lattice possesses features that are distinct from those formed by other membrane-bending proteins. Whereas endophilin ^27^ and dynamin ^30^ assemble as 2-start helical arrays, and SNX1 ^29^ and ACAP1 ^28^ assemble as 1-start helical arrays, we find that ARF6 assembles as 5-start arrays. Another notable distinction is that ARF6 forms tetramers as the basic building block of its lattice structure, while dimers are involved for lattices formed by endophilin ^27^, SNX1 ^29^ and dynamin ^30^, and a dimer of dimer is involved in the ACAP1 lattice ^28^.

Previous studies on vesicle formation by two of the best characterized coat complexes have revealed that the ability of ARF1 and Sar1 to induce liposome tubulation reflects their roles in COPI and COPII vesicle fission, respectively ^19,20^. We have now advanced a molecular understanding of this role through the structural elucidation of the ARF lattice assembled on tubulated membrane. Central to concluding that this structure explains how ARF acts in vesicle fission has been functional studies that involve mutating key residues predicted to be critical for lattice assembly and then finding that they inhibit COPI vesicle fission.

We have also ruled out a key caveat of this approach, which is that the mutations could be disrupting instead ARF1 interacting with other proteins that act in COPI vesicle formation. Initially, we compared the positioning of ARF1 in the lattice structure to that in complex with coatomer on COPI vesicles, which had been previously elucidated by cryo-EM ^15^. The results predicted that the mutations targeting the major protein interfaces of the ARF1 lattice are unlikely to have a significant impact on the ability of ARF1 to assemble with coatomer for vesicle formation. As further support, we noted that the mutations allow vesicle formation to proceed through the budding stage in which the ARF1-coatomer interactions should play a central role. We also performed functional studies for further confirmation, which interrogated ARF1 interacting with coatomer, its regulators for proper recruitment to the Golgi, as well as ARFGAP and BARS. The results further supported that the mutations have specific effects in disrupting the ARF1 lattice structure in explaining how COPI vesicle fission is inhibited.

Our studies on the ARF1 lattice have also shed new insights into how COPI vesicle fission is achieved. As ARF1, ARFGAP1, and BARS have all been identified to act in this process ^20,33,41^, a major mechanistic question has been whether their roles would involve these proteins acting simultaneously at the bud neck, or could they act more distinctly. Our finding that COPI vesicle fission involves the existence of an ARF1 lattice structure now points to the latter explanation as being more likely.

We have also performed additional studies that further support, and expand on, this conclusion. Our pursuit of the COPI vesicle reconstitution system reveals that mutations which disrupt the ARF1 lattice arrest COPI vesicle formation starting in early fission. Thus, when also considering that a previous study has found that inhibiting the GAP activity results in COPI buds having severely constricted neck, reflecting a block in late fission ^33^, the collective observations point to the action of the ARF lattice preceding that of the GAP activity during vesicle fission. Further supporting this conclusion, we find that the ARF lattice allows the GAP to dock onto the structure to catalyze ARF-GTP hydrolysis. Functionally, we also observe vesiculation of ARF6 tubules when a GAP (ACAP1) is added to these tubules.

Vesicle formation involves three major sequential stages: i) coat recruitment, ii) vesicle budding, and iii) vesicle fission. Studies over the years have revealed that ARF proteins play key roles in all three stages. For the first two stages, structural studies have achieved a molecular understanding of how ARF acts ^11,13–15^. Our structural elucidation of an ARF-induced tubular lattice has now achieved a molecular understanding of how ARF acts in the third stage (summarized in **Extended Data Fig. 7**).

It is notable that GTP-binding by dynamin has been found to form a tubular lattice structure around the clathrin bud neck, followed by GTP hydrolysis that results in neck scission to complete vesicle formation ^42,43^. In this regard, our elucidation that the ARF tubular lattice is involved in vesicle fission and this structure also allows docking by the GAP for the subsequent vesiculation of the ARF-induced tubules has the key implication that mechanisms of vesicle fission are fundamentally conserved.

## METHODS

### ARF6 protein preparation

Human ARF6 (NP_001654.1) was subcloned into the pET22b vector, co-transformed with N-myristoyltransferase in the pBB131 vector, and co-expressed as fusion proteins with C-terminal 6xHis tag in Escherichia coli BL21 (DE3) cells. Cells were grown in NZCYM medium at 27°C until the OD at 600 nm reached to ∼0.4, then the myristate solution is added. When OD_600 nm_ reached to ∼1.2, cells were induced at 27°C for 5∼6 hours with 0.5 mM IPTG. Cells were harvested by centrifugation, re-suspended in lysis buffer containing 50 mM Tris pH 8.0, 300 mM NaCl, 1 mM MgCl_2_, 0.5 mM dithiothreitol (DTT), 0.5% Triton X-100, 10 mM imidazole, and then disrupted by a high-pressure homogenizer (JINBIO). After centrifugation for 40 minutes at 16,000 rpm, the supernatant was collected and incubated with Ni-NTA beads (Roche) at 4°C, washed with buffer containing 50 mM Tris pH 8.0, 300 mM NaCl, 1 mM MgCl_2_, 30 mM imidazole, and then eluted with 50 mM Tris pH 8.0, 300 mM NaCl, 1 mM MgCl_2_, 400 mM imidazole. The eluted protein was brought slowly to 35% saturation (19.4g/100ml) in ammonium sulfate. After stirring on ice for 30 minutes, the protein mixture was centrifuged at 12,000 rpm for 30 minutes. The precipitate was dissolved into buffer containing 50mM Hepes pH 7.4,100mM NaCl and 1mM MgCl_2_, and then centrifuged at 16,000 rpm for 40 minutes. The supernatant was collected and then stored at −80°C.

To generate mutant forms of ARF6, site-directed mutagenesis of select residues was performed by overlap PCR. Expression and purification of ARF6 mutants were done as described above for the wild-type form.

For nucleotide loading, ARF6 was incubated with GTP or GDP in a nucleotide exchange buffer (50 mM HEPES-KOH pH7.4, 100 mM NaCl, 1 mM MgCl_2_, 2 mM EDTA and 1mM GTP or GDP) at 30 °C for 30 minutes. Reaction was then terminated by increasing the MgCl_2_ concentration (5 mM final) to stabilize the nucleotide bound state.

### Liposome assays

All lipids were purchased from Avanti Polar lipids. Lipid mixtures, containing 40% phosphatidylcholine (DOPC), 30% phosphatidylethanolamine (DOPE), 20% phosphatidylserine, 8% phosphatidylinositol-4,5-bisphosphate [PI(4,5)P2] and 2% cholesterol, were dried under nitrogen gas and then kept under vacuum for at least three hours. Dried lipid mixtures were resuspended in 50 mM HEPES, pH7.4, 50 mM NaCl for 30 minutes at 37°C, and then underwent freeze-thaw (from liquid nitrogen to 37°C) for 5 cycles, followed by extrusion through membrane filters of 0.2 um to generate liposomes of 200 nm diameter.

For liposome tubulation analysis, ARF6 protein (0.4mg/ml) was incubated with liposomes (0.5mg/ml) at room temperature. After 30 minutes, the incubation mixture was applied onto a glow-discharged carbon-coated EM grid and stained with uranyl acetate. EM grids were examined with a transmission electron microscope (FEI Tecnai Spirit 120) and the micrographs were recorded under the nominate magnification of 49,000X.

### Cryo-electron microscopy

ARF6 protein (0.8mg/ml) was incubated with 200 nm-diameter liposomes (1.7 mg/ml) at room temperature for 30 minutes. A drop (3.0 µl) of the mixture was applied onto a Quantifoil 300-mesh R2/1 holy carbon grid that was pre-treated in plasma cleaner. The grid was then blotted for 4 seconds with a blot force 0 at 100% humidity, using FEI Vitrobot (Mark IV), before it was quickly frozen in liquid ethane that was cooled by liquid nitrogen.

ARF6 coated membrane tubules were imaged with an FEI Titan Krios cryo-electron microscope that was operated under 300kV and equipped with a direct electron detector device Falcon III camera (Thermo Fisher Scientific Inc.). Low-dose images (∼50e^-^/Å^2^) were collected manually. The nominal magnification was set to 75,000X, which corresponds to a pixel size 1.1 Å. The defocus range was set to 1.5∼2.2 µm. A total of 1366 cryo-EM micrograph’s movie stacks were collected. Motion correction and defocus estimation for all micrographs were performed using MotionCorr2 ^44^ and GCTF ^45^ respectively. 1044 ARF6-coated tubules were boxed using e2helixboxer.py in the package of EMAN2 ^46^ with a box width of 384 pixels. Most of the tubules had similar diameters, as assessed by the diameter measurement of peak distance of the 1-dimensional projection of tubules (**Extended Data Fig. 2**). We further checked the diffraction patterns of tubules by randomly selecting 50 out of 1104 total. The diffraction patterns were homogeneous as assessed manually.

### Helical reconstruction

With the consistency of both the diameter and diffraction pattern, we averaged all the diffraction patterns together. We then indexed the average pattern to calculate all possible helical parameters for the symmetry. All 1104 tubules were segmented into particles and a stack with 56557 particles were generated with boxsize 384. In order to quickly screen for the correct helical parameters, about half of the particle stack with 27430 particles were shrunken 4 times and used to perform iterative helical real space reconstruction (IHRSR) ^47^. Cylinder with 34 nm diameters was generated by Spider ^48^ and set as the initial model for IHRSR reconstructions ^22^. After dozens of IHRSR reconstructions, we found that a helical rise of 27.8 Å and twist 27.5°, was the best one, because its density shape was similar to that of the ARF6 crystal structure. We then refined this structure using IHRSR with the raw scale 384, and imposed the dihedral symmetry present in the asymmetric units. These additional steps further improved the resolution (**Extended Data Fig. 1d**). The diffraction pattern of projections from the map was also consistent to that of raw tubules (**Extended Data Fig. 1d**), which confirmed that the indexing was correct.

We then used RELION to refine the structure. Initially, the whole dataset of 56,557 particles were sorted by 2D class averaging. Subsequently, 56,407 particles were analyzed using 3D refinement, with the initial parameters of helical rise 27.97 Å and twist 27.28° and the initial 3D model from IHRSR reconstruction. This step yielded a cryo-EM map with the resolution of 4.2 Å. After CTF refinement and particle polishing, the resolution was extended to 4.1 Å. We then performed further 3D classification to sort particles and selected classes 2 and 7 that have the closest helical parameters for the last round of 3D refinement, which yielded the final map with a resolution of 3.9 Å according to the FSC (Fourier Shell Correlation) gold standard. The final refined helical parameters from RELION 3D refinement were reported as 28.01 Å for helical rise and 27.28° for helical twist. We then performed density modification using PHENIX {Adams, 2010 #30}, resulting in a map with an estimated resolution of 3.6 Å. This map was used for the subsequent model building.

### Model building and refinement

We used the crystal structure of GTPγS-bound ARF6 (PDB ID: 2J5X) as an initial template to dock into the cryo-EM map by rigid body fitting. The model was manually edited in Chimera ^24^ or Chimera X, and a new model with four polypeptide chains was generated as the asymmetric tetramer unit. This tetramer was refined in Rosetta ^49^ with helical symmetry. The final model was selected according to the validation-map agreement, model geometry and full-map agreement. The model was further refined in Phenix ^50^ to improve the stereochemistry.

### Molecular dynamics simulation

ARF6 tetramer structure was extracted from our cryo-EM structure, while a myristoylated N-terminal fragment of Arf6 (PDB:2BAO) was added to the N-terminus region. The protonation states of histidine residues were determined by local hydrogen-bonding interactions. A symmetry lipid bilayer with 452 lipids was generated using CHARMM-GUI Membrane Builder ^51,52^. The membrane is composed of 180 molecules of DOPC, 136 of DOPE, 90 of POPS, 36 of PIP2 and 10 of cholesterol. The membrane was equilibrated for 100ns and then merged with the protein. The protein was placed at least 1.0 nm above the upper leaflet of the membrane. The membrane and protein-membrane systems were solvated with TIP3P^53^ water molecules in a periodic box, and were then neutralized by adding 150mM of sodium and chloride ions. A system consisting of only the protein and solution was also modeled.

All simulations were conducted by GROMACS-2020^54^ along with CHARMM36 force field^55,56^. The system was energy minimized for 10,000 steps with 1000 KJ·mol^-1^·nm^-2^ harmonic restraints on the protein backbone and lipids, then the restrained systems were heated to 310K over a period of 2.5ns under constant temperature/pressure (*NPT*) conditions. This was followed by two equilibration simulations of 2.5 ns each, with gradually decreased restraints. The production simulation was executed for 800ns. An integration time step of 2 fs was applied while all hydrogen-containing covalent bonds were constrained by the LINCS algorithm^57^. The pressure of the models was maintained at 1 bar by a semi-isotropic Parrinello-Rahman barostat^58^, while the temperature was controlled by Nose−Hoover thermostat^59^. The particle-mesh Ewald method^60^ was employed to calculate long-range electrostatic interactions, whereas a smooth (10-12 Å) cutoff was applied to treat van der Waals (vdW) interactions.

Model visualization and representation were performed in Visual Molecular Dynamics (VMD) software^61^. Analysis was conducted using locally written scripts and built-in tools in Gromacs.

### TfR recycling assay

TfR transport from the recycling endosome to the plasma membrane was performed essentially as previously described ^10^. Briefly, HeLa cells were incubated with Alexa 546-conjugated Tf (5μg/ml in DMEM) at 37°C for 2 hours to allow the steady-state accumulation of Tf in endosomes. Subsequently, cells were incubated with medium without Tf for different times as indicated in the figures. Cells were then stained for cellubrevin (a marker for the recycling endosome), followed by confocal microscopy to assess the exit of Tf from the recycling endosome.

### COPI transport assay

COPI transport from the Golgi to the ER was performed essentially as previously described ^38^. In brief, HeLa cells were transfected with VSVG-ts045-KDELR-Myc in the pRose vector for 1 day, and then incubated at 32°C for 8 hours to achieve steady-state distribution at the Golgi. Cells were then shifted to 40°C for different times as indicated in the figures. Cells were then stained for giantin, followed by confocal microscopy to assess the exit of VSVG-ts045-KDELR from the Golgi.

### In vitro reconstitution of COPI vesicle formation

The preparation of recombinant forms of ARF1, ARFGAP1 BARS, and the purification of coatomer from liver tissue have been described ^62^. The isolation of Golgi membrane from cells has also been described ^63^. The incubation of these components for the reconstitution of COPI vesicles was done as described previously ^62^, with minor modification.

Briefly, salt-washed (3M KCl) Golgi membrane (100 µg) was pelleted by centrifugation at 17,000 x g at 4°C for 30 minutes. The pellet containing Golgi membrane was resuspended using 1 ml of washing buffer (25 mM HEPES-KOH pH 7.4, 3M KCl, 2.5 mM Mg(OAc)_2_, 1 mg/ml soybean trypsin inhibitor), followed by incubation on ice/water for 5 minutes, and then incubated with myristoylated ARF1 (6 μg/ml), ARFGAP1 (6 μg/ml), BARS (3 μg/ml), coatomer (6 μg/ml), and GTP (2 mM) at 37°C for 30 minutes in 300 μl of reaction buffer (25 mM HEPES-KOH pH 7.4, 50 mM KCl, 2.5 mM Mg(OAc)_2_, 1 mg/ml soybean trypsin inhibitor, 1 mg/ml BSA, and 200 mM sucrose). After the incubation, centrifugation was performed at 17,000 x g at 4°C for 30 minutes to pellet Golgi membrane. Centrifugation was then performed on the supernatant at 200,000 x g using TLA120.2 rotor at 4°C for 30 minutes to pellet vesicles. The level of reconstituted COPI vesicles was determined by western blotting using antibody against β-COP.

EM examination of Golgi membrane using the whole-mount technique has been described previously^38^. Briefly, the sample was placed on carbon grid (CF200-CU, Electron Microscopy Science). The carbon grid was washed twice with water twice and stained using 1% uranyl acetate. Golgi membranes were then examined using a JEOL JEM-1011 electron microscope. Quantitation involved the examination of 10 meshes per condition.

### Golgi membrane recruitment

The Golgi membrane (40 µg) was pelleted by centrifugation at 17,000 x g at 4°C for 30 minutes. The pelleted membrane was then resuspended using 100 µl of reaction buffer (25 mM HEPES-KOH pH 7.4, 50 mM KCl, 2.5 mM Mg(OAc)_2_, and 200 mM sucrose) and incubated with GTP or GMPPNP loaded ARF1 (2 µg) and coatomer (6 µg) at 37°C for 30 minutes. After the incubation, centrifugation was performed at 17,000 x g at 4°C for 30 minutes to pellet Golgi membrane. The level of ARF1 or COPI recruitment on Golgi membrane was determined by western blotting using antibodies against β-COP or ARF1.

### Assessing Golgi localization of ARF1

The ARF1 WT and mutants (M1∼M5) were subcloned into the pcDNA 3.1 vector. Hela cells were transfected with si-ARF1 (5’-AACATCTTCGCCAACCTCTTC-3’) for 1 day and then ARF1 WT and mutants were expressed for 1 day. Cells were then stained using antibodies against giantin (1:500) and ARF1 (1:500). Cells were imaged using Zeiss LSM900 confocal microscopy and Zen 2.3 confocal acquisition software. The colocalization of ARF1 with Giantin was measured using MetaMorph 7.7 (MDS Analytical Technologies).

### GST pulldown

GST fusion proteins were expressed in bacteria (BL21) by IPTG induction. The cells were collected by centrifugation 4,000 x g at 4 °C for 10 minutes and then lysed using lysis buffer (20 mM Tris-HCl pH 7.4, 100 mM NaCl, 1% Triton X-100 and protease inhibitor) at 4 °C for 30 minutes. The cell lysates were centrifuged at 4,000 x g at 4 °C for 15 minutes. The clear supernatant fraction was incubated with glutathione Sepharose bead (GE healthcare). GST fusion proteins on beads were incubated with recombinant proteins at 25 °C for 2 hours or 4 °C overnight in reaction buffer (25 mM Tris-HCl pH 7.4, 50 mM KCl, 2.5 mM Mg(OAc)_2_, and 0.1 % Triton X-100). Beads were washed twice using 1 ml of reaction buffer, and the bound proteins were detected by western blotting.

## DATA AVAILABILITY

The EM map of helical tubules has been deposited in Electron Microscopy Data Bank (EMDB) with the accession code EMD-33414. The corresponding coordinates of ARF6 helical array on the tubule has been deposited in the PDB with the accession code 7XRD.

## AUTHOR CONTRIBUTIONS

F.S., S.Y.P. and V.W.H. supervised the project. X.P., S.Y.P., V.W.H. and F.S. designed experiments. X. P. and Y. Z. performed EM work. X.P. performed purification experiments and mutagenesis studies with the help of J.X., G.Y., T.Z. and Y.W.. J.L. performed the TfR recycling assay. K.P. and H.M.T performed the COPI transport assay and the COPI vesicle reconstitution system under the supervision of S.Y.P.. Z.L. performed MD simulation under the supervision of J.F.. X.P., Y.Z., H.M.T., Z.L., J.L., E.H.E., J.F., S.Y.P., V.H., and F.S. analyzed the data. All contributed to manuscript preparation. X.P., S.Y.P, V.H. and F.S. wrote the manuscript.

## ACKNOWLEDGMENTS

This work was supported by grants from the National Natural Science Foundation of China (31961160723, 31670744, 31770794 and 31925026) to F.S., X.P. and Y.Z., the National Institutes of Health USA (R37GM058615 to V.W.H. and R35GM122510 to E.H.E.), the National Research Foundation of Korea (2021R1A6A1A10042944, 2017R1A5A1015366, 2020R1C1C1008823) to S.Y.P., and NSFC/RGC Joint Research Scheme N_CityU104/19 and Hong Kong Research Grant Council Collaborative Research Fund: C6021-19E to J.F..

We would like to thank Ping Shan, Ruigang Su and Mengyue Lou for assistance in lab management, and Hanlin Wang for assistance in experiments. All EM data were collected at Center for Biological Imaging (CBI, http://cbi.ibp.ac.cn), Institute of Biophysics, Chinese Academy of Sciences. We are grateful to Dr. Xiaojun Huang and Boling Zhu (CBI) for their assistance in EM data collection.

## REFERENCES

1 D’Souza-Schorey, C. & Chavrier, P. ARF proteins: roles in membrane traffic and beyond. Nat Rev Mol Cell Biol 7, 347–358 (2006).

2 Donaldson, J. G. & Jackson, C. L. ARF family G proteins and their regulators: roles in membrane transport, development and disease. Nat Rev Mol Cell Biol 12, 362–375 (2011).

3 Donaldson, J. G., Cassel, D., Kahn, R. A. & Klausner, R. D. ADP-ribosylation factor, a small GTP-binding protein, is required for binding of the coatomer protein beta-COP to Golgi membranes. Proc Natl Acad Sci USA 89, 6408–6412 (1992).

4 Serafini, T. et al. ADP-ribosylation factor is a subunit of the coat of Golgi-derived COP-coated vesicles: a novel role for a GTP-binding protein. Cell 67, 239–253 (1991).

5 Traub, L. M., Ostrom, J. A. & Kornfeld, S. Biochemical dissection of AP-1 recruitment onto Golgi membranes. J Cell Biol 123, 561–573 (1993).

6 Ooi, C. E., Dell’Angelica, E. C. & Bonifacino, J. S. ADP-Ribosylation factor 1 (ARF1) regulates recruitment of the AP-3 adaptor complex to membranes. J Cell Biol 142, 391–402 (1998).

7 Boehm, M., Aguilar, R. C. & Bonifacino, J. S. Functional and physical interactions of the adaptor protein complex AP-4 with ADP-ribosylation factors (ARFs). Embo J 20, 6265–6276 (2001).

8 Barlowe, C. et al. COPII: a membrane coat formed by Sec proteins that drive vesicle budding from the endoplasmic reticulum. Cell 77, 895–907 (1994).

9 Li, J. et al. An ACAP1-containing clathrin coat complex for endocytic recycling. J Cell Biol 178, 453–464, doi:10.1083/jcb.200608033 (2007).

10 Hsu, J. W. et al. The protein kinase Akt acts as a coat adaptor in endocytic recycling. Nat Cell Biol 22, 927–933 (2020).

11 Yu, X., Breitman, M. & Goldberg, J. A structure-based mechanism for Arf1-dependent recruitment of coatomer to membranes. Cell 148, 530–542, doi:10.1016/j.cell.2012.01.015 (2012).

12 Bi, X., Corpina, R. A. & Goldberg, J. Structure of the Sec23/24-Sar1 pre-budding complex of the COPII vesicle coat. Nature 419, 271–277 (2002).

13 Faini, M. et al. The structures of COPI-coated vesicles reveal alternate coatomer conformations and interactions. Science 336, 1451–1454, doi:10.1126/science.1221443 (2012).

14 Dodonova, S. O. et al. VESICULAR TRANSPORT. A structure of the COPI coat and the role of coat proteins in membrane vesicle assembly. Science 349, 195–198, doi:10.1126/science.aab1121 (2015).

15 Dodonova, S. O. et al. 9A structure of the COPI coat reveals that the Arf1 GTPase occupies two contrasting molecular environments. Elife 6, doi:10.7554/eLife.26691 (2017).

16 Zanetti, G. et al. The structure of the COPII transport-vesicle coat assembled on membranes. Elife 2, e00951, doi:10.7554/eLife.00951 (2013).

17 Hutchings, J., Stancheva, V., Miller, E. A. & Zanetti, G. Subtomogram averaging of COPII assemblies reveals how coat organization dictates membrane shape. Nat Commun 9, 4154, doi:10.1038/s41467-018-06577-4 (2018).

18 Hutchings, J. et al. Structure of the complete, membrane-assembled COPII coat reveals a complex interaction network. Nat Commun 12, 2034, doi:10.1038/s41467-021-22110-6 (2021).

19 Lee, M. C. et al. Sar1p N-terminal helix initiates membrane curvature and completes the fission of a COPII vesicle. Cell 122, 605–617, doi:10.1016/j.cell.2005.07.025 (2005).

20 Beck, R. et al. Coatomer and dimeric ADP ribosylation factor 1 promote distinct steps in membrane scission. J Cell Biol 194, 765–777, doi:10.1083/jcb.201011027 (2011).

21 Lundmark, R., Doherty, G. J., Vallis, Y., Peter, B. J. & McMahon, H. T. Arf family GTP loading is activated by, and generates, positive membrane curvature. Biochem J 414, 189–194, doi:10.1042/BJ20081237 (2008).

22 Egelman, E. H. The iterative helical real space reconstruction method: surmounting the problems posed by real polymers. Journal of structural biology 157, 83–94, doi:10.1016/j.jsb.2006.05.015 (2007).

23 Zivanov, J. et al. New tools for automated high-resolution cryo-EM structure determination in RELION-3. eLife 7, doi:10.7554/eLife.42166 (2018).

24 Pettersen, E. F. et al. UCSF chimera - A visualization system for exploratory research and analysis. J Comput Chem 25, 1605–1612, doi:10.1002/jcc.20084 (2004).

25 Pasqualato, S., Menetrey, J., Franco, M. & Cherfils, J. The structural GDP/GTP cycle of human Arf6. EMBO Rep 2, 234–238, doi:10.1093/embo-reports/kve043 (2001).

26 Menetrey, J., Macia, E., Pasqualato, S., Franco, M. & Cherfils, J. Structure of Arf6-GDP suggests a basis for guanine nucleotide exchange factors specificity. Nature structural biology 7, 466–469 (2000).

27 Mim, C. et al. Structural basis of membrane bending by the N-BAR protein endophilin. Cell 149, 137–145, doi:10.1016/j.cell.2012.01.048 (2012).

28 Pang, X. et al. A PH Domain in ACAP1 Possesses Key Features of the BAR Domain in Promoting Membrane Curvature. Dev Cell 31, 73–86, doi:10.1016/j.devcel.2014.08.020 (2014).

29 Zhang, Y. et al. Structural insights into membrane remodeling by SNX1. Proc Natl Acad Sci U S A 118, e2022614118, doi:10.1073/pnas.2022614118 (2021).

30 Kong, L. et al. Cryo-EM of the dynamin polymer assembled on lipid membrane. Nature 560, 258–262, doi:10.1038/s41586-018-0378-6 (2018).

31 Krauss, M. et al. Arf1-GTP-induced tubule formation suggests a function of Arf family proteins in curvature acquisition at sites of vesicle budding. J Biol Chem 283, 27717–27723 (2008).

32 Bielli, A. et al. Regulation of Sar1 NH2 terminus by GTP binding and hydrolysis promotes membrane deformation to control COPII vesicle fission. J Cell Biol 171, 919–924 (2005).

33 Yang, J. S. et al. GAPDH inhibits intracellular pathways during starvation for cellular energy homeostasis. Nature 561, 263–267, doi:10.1038/s41586-018-0475-6 (2018).

34 Ismail, S. A., Vetter, I. R., Sot, B. & Wittinghofer, A. The structure of an Arf-ArfGAP complex reveals a Ca2+ regulatory mechanism. Cell 141, 812–821 (2010).

35 Serafini, T. et al. A coat subunit of Golgi-derived non-clathrin-coated vesicles with homology to the clathrin-coated vesicle coat protein beta-adaptin. Nature 349, 215–220 (1991).

36 Yang, J. S. et al. ARFGAP1 promotes the formation of COPI vesicles, suggesting function as a component of the coat. J Cell Biol 159, 69–78 (2002).

37 Shiba, T. et al. Molecular mechanism of membrane recruitment of GGA by ARF in lysosomal protein transport. Nat Struct Biol 10, 386–393, doi:10.1038/nsb920 (2003).

38 Park, S. Y. et al. The late stage of COPI vesicle fission requires short forms of phosphatidic acid and diacylglcerol. Nature communications Jul 30; 10(1), 3409, doi:10.1038/s41467-019-11324-4 (2019).

39 Yang, J. S. et al. COPI acts in both vesicular and tubular transport. Nat Cell Biol 13, 996–1003 (2011).

40 Faini, M., Beck, R., Wieland, F. T. & Briggs, J. A. Vesicle coats: structure, function, and general principles of assembly. Trends Cell Biol 23, 279–288, doi:10.1016/j.tcb.2013.01.005 (2013).

41 Yang, J. S. et al. A role for BARS at the fission step of COPI vesicle formation from Golgi membrane. EMBO J 24, 4133–4143 (2005).

42 Takei, K., McPherson, P. S., Schmid, S. L. & De Camilli, P. Tubular membrane invaginations coated by dynamin rings are induced by GTP-gamma S in nerve terminals. Nature 374, 186–190 (1995).

43 Mattila, J. P. et al. A hemi-fission intermediate links two mechanistically distinct stages of membrane fission. Nature 524, 109–113, doi:10.1038/nature14509 (2015).

44 Zheng, S. Q. et al. MotionCor2: anisotropic correction of beam-induced motion for improved cryo-electron microscopy. Nature methods 14, 331–332, doi:10.1038/nmeth.4193 (2017).

45 Zhang, K. Gctf: Real-time CTF determination and correction. Journal of structural biology 193, 1–12, doi:10.1016/j.jsb.2015.11.003 (2016).

46 Tang, G. et al. EMAN2: an extensible image processing suite for electron microscopy. Journal of structural biology 157, 38–46, doi:10.1016/j.jsb.2006.05.009 (2007).

47 Egelman, E. H. Reconstruction of Helical Filaments and Tubes. Method Enzymol 482, 167–183, doi:10.1016/S0076-6879(10)82006-3 (2010).

48 Shaikh, T. R. et al. SPIDER image processing for single-particle reconstruction of biological macromolecules from electron micrographs. Nat Protoc 3, 1941–1974, doi:10.1038/nprot.2008.156 (2008).

49 Wang, R. Y. et al. Automated structure refinement of macromolecular assemblies from cryo-EM maps using Rosetta. eLife 5, doi:10.7554/eLife.17219 (2016).

50 Adams, P. D. et al. PHENIX: a comprehensive Python-based system for macromolecular structure solution. Acta Crystallogr D Biol Crystallogr 66, 213–221, doi:10.1107/S0907444909052925 (2010).

51 Jo, S., Kim, T., Iyer, V. G. & Im, W. CHARMM-GUI: a web-based graphical user interface for CHARMM. J Comput Chem 29, 1859–1865, doi:10.1002/jcc.20945 (2008).

52 Lee, J. et al. CHARMM-GUI Input Generator for NAMD, GROMACS, AMBER, OpenMM, and CHARMM/OpenMM Simulations Using the CHARMM36 Additive Force Field. J Chem Theory Comput 12, 405–413, doi:10.1021/acs.jctc.5b00935 (2016).

53 Jorgensen, W. L., Chandrasekhar, J., Madura, J. D., Impey, R. W. & Klein, M. L. Comparison of Simple Potential Functions for Simulating Liquid Water. J Chem Phys 79, 926–935, doi:Doi 10.1063/1.445869 (1983).

54 Abraham M. J., M. T., Schulz R., Páll S., Smith J. C., Hess B. and Lindahl E. GROMACS: High performance molecular simulations through multi-level parallelism from laptops to supercomputers. SoftwareX 1, 19–25, doi:10.1016/j.softx.2015.06.001 (2015).

55 Best, R. B. et al. Optimization of the Additive CHARMM All-Atom Protein Force Field Targeting Improved Sampling of the Backbone φ, ψ and Side-Chain χ and χ Dihedral Angles. Journal of Chemical Theory and Computation 8, 3257–3273, doi:10.1021/ct300400x (2012).

56 Klauda, J. B. et al. Update of the CHARMM all-atom additive force field for lipids: validation on six lipid types. J Phys Chem B 114, 7830–7843, doi:10.1021/jp101759q (2010).

57 Hess, B., Bekker, H., Berendsen, H. J. C. & Fraaije, J. G. E. M. LINCS: A linear constraint solver for molecular simulations. J Comput Chem 18, 1463–1472, doi:Doi 10.1002/(Sici)1096-987x(199709)18:12<1463::Aid-Jcc4>3.0.Co;2-H (1997).

58 Parrinello, M. & Rahman, A. Polymorphic Transitions in Single-Crystals - a New Molecular-Dynamics Method. J Appl Phys 52, 7182–7190, doi:Doi 10.1063/1.328693 (1981).

59 Nose, S. A Molecular-Dynamics Method for Simulations in the Canonical Ensemble. Mol Phys 52, 255–268, doi:Doi 10.1080/00268978400101201 (1984).

60 Darden, T., York, D. & Pedersen, L. Particle Mesh Ewald - an N.Log(N) Method for Ewald Sums in Large Systems. J Chem Phys 98, 10089–10092, doi:Doi 10.1063/1.464397 (1993).

61 Humphrey, W., Dalke, A. & Schulten, K. VMD: Visual molecular dynamics. J Mol Graph Model 14, 33-38, doi:Doi 10.1016/0263-7855(96)00018-5 (1996).

62 Park, S. Y., Yang, J. S. & Hsu, V. W. Reconstitution of COPI Vesicle and Tubule Formation. Methods Mol Biol 1496, 63–74 (2016).

63 Fowler, P. W. et al. Membrane stiffness is modified by integral membrane proteins. Soft matter 12, 7792–7803 (2016).

