## Extended Data for "Structural elucidation of how ARF small GTPases induce membrane tubulation for vesicle fission"

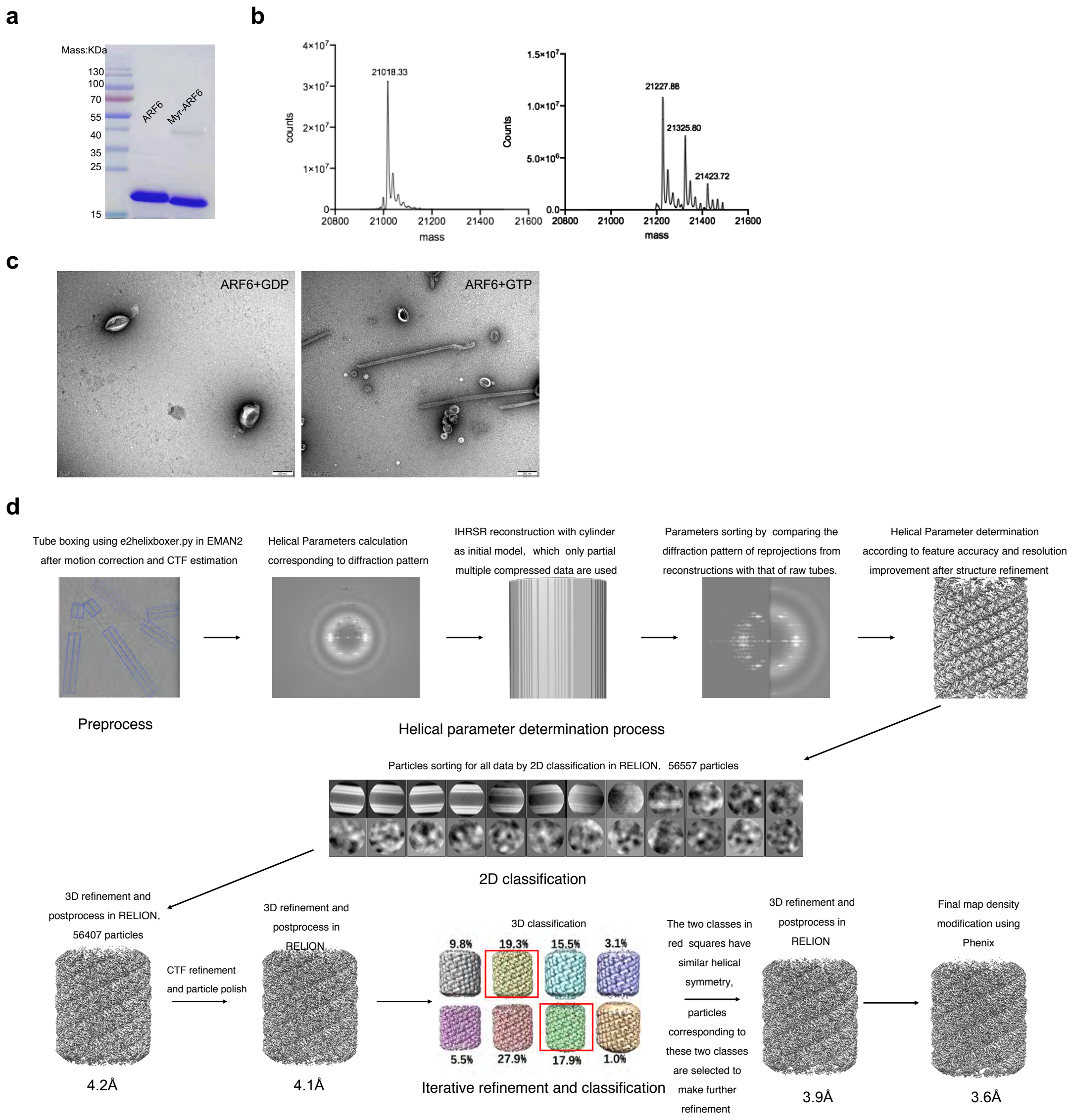

**Extended Data Fig. 1. Characterizing recombinant ARF6 and summary of cryo-EM processing**

**a,** Gel analysis of purified ARF6 by Coomassie staining, comparing myristoylated and unmyristoylated forms. **b**, Mass spectrometry analysis comparing ARF6 in unmyristoylated (left) versus myristoylated (right) form. The molecular mass difference of 209.6 Da is consistent with the addition of a myristoyl group of 210.2 Da. **c**, EM examination assessing tubulation activity by ARF6 in its nucleotide-bound states as indicated. Scale bar, 200 nm. **d,** Workflow of cryo-EM data processing. See Methods section for a detailed description of the different steps.

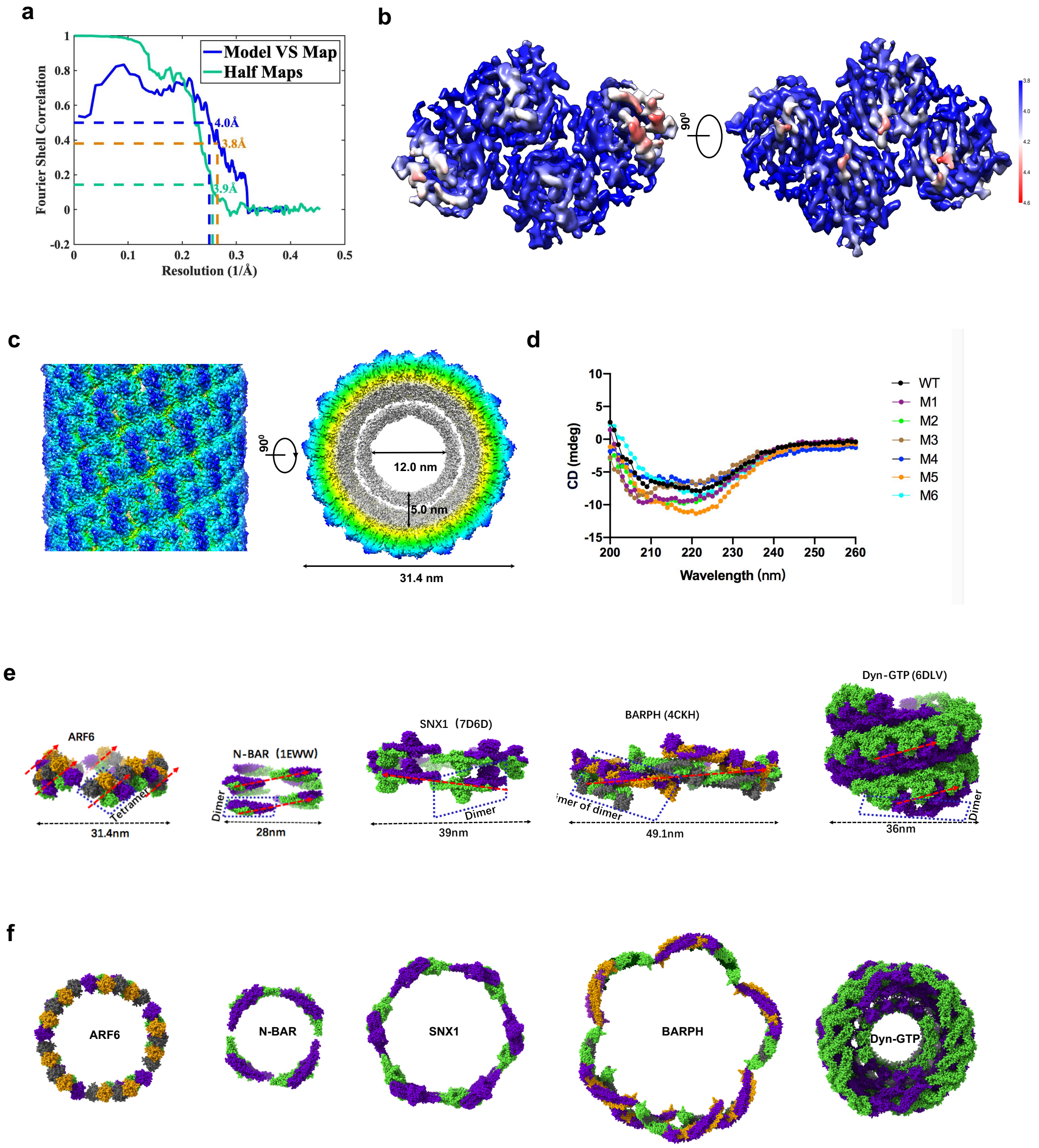

**Extended Data Fig. 2. Further characterization of the ARF6 lattice.**

**a**, FSC (Fourier shell correlation) curve from two independently refined datasets indicates overall map resolution of 3.9 Å (FSC = 0.143) (green curve). The resolution, validated by the cross-FSC between cryo-EM map and generated model (blue curve), is 4.0 Å at FSC = 0.5, and 3.8 Å at FSC=0.38 (orange lines; 0.38 is the square root of 0.143). Since the structural model does not contain membrane information, the FCS curve of model-to-map is worse at the low resolution region in comparison with that of half maps. **b**, Local resolution map is depicted as an isosurface, colored according to the resolution. **c**, Cryo-EM map of the ARF6-coated tubule, with the side view (left) showing the surface of the ARF6 lattice, and the cross-sectional view (right) showing the diameters of the inner and outer surface of the membrane and the depth of the lattice coating the membrane. The map is colored according to the radius. **d**, Circular dichroism spectroscopy of ARF6 wild type and mutants. **e,** The basic (asymmetric) unit of each helical array is indicated by the blue parallelogram. The asymmetric unit of ARF6 assembly is a tetramer. Each subunit is shown in a different color: gold, grey, green and purple. The asymmetric unit of assembly by SNX1, N-BAR or dynamin1 is the dimer, with subunits colored in green and purple. The asymmetric unit of ACAP1 (BAR-PH domain) assembly is a dimer of dimer, colored in gold, purple, green and grey. The red arrow indicates the direction of helical start. **f,** Cross-sectional views of the helical assemblies of ARF6, N-BAR, SNX1, ACAP1 (BARPH domain) and dynamin on membrane tubules.

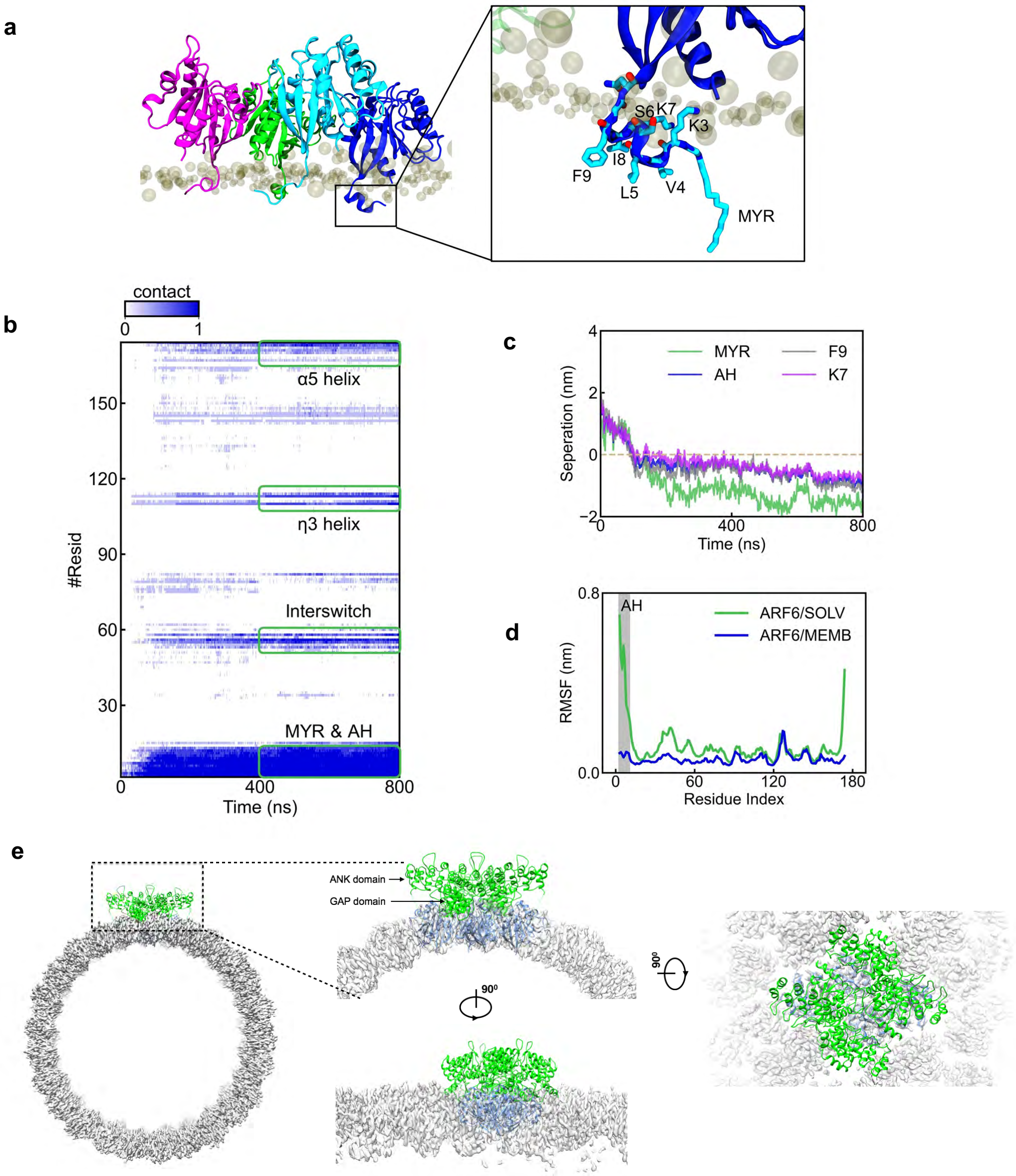

**Extended Data Fig. 3. Membrane insertion by the ARF6 tetramer and GAP docking onto the ARF6 lattice.**

**a**, Snapshot of ARF6 tetramer on the membrane at the final frame of the molecular dynamics simulation, and zoomed-in view is focusing on an N-terminal myristoyl chain (MYR) and an amphipathic helix (AH) engaging the membrane. **b**, Time course of protein contacts with lipids. **c**, Time evolved profile of the separation between the lipid membrane and MYR, AH, F9 and K7. **d**, Root-mean-square-fluctuation (RMSF) profiles of ARF6 in solution and on the membrane. **e**,

Superposition of ARF6-ASAP3 complex structure onto the ARF6 lattice structure, showing that GAP and ANK domains of ASAP3 can dock onto the surface of the ARF lattice without structural collision.

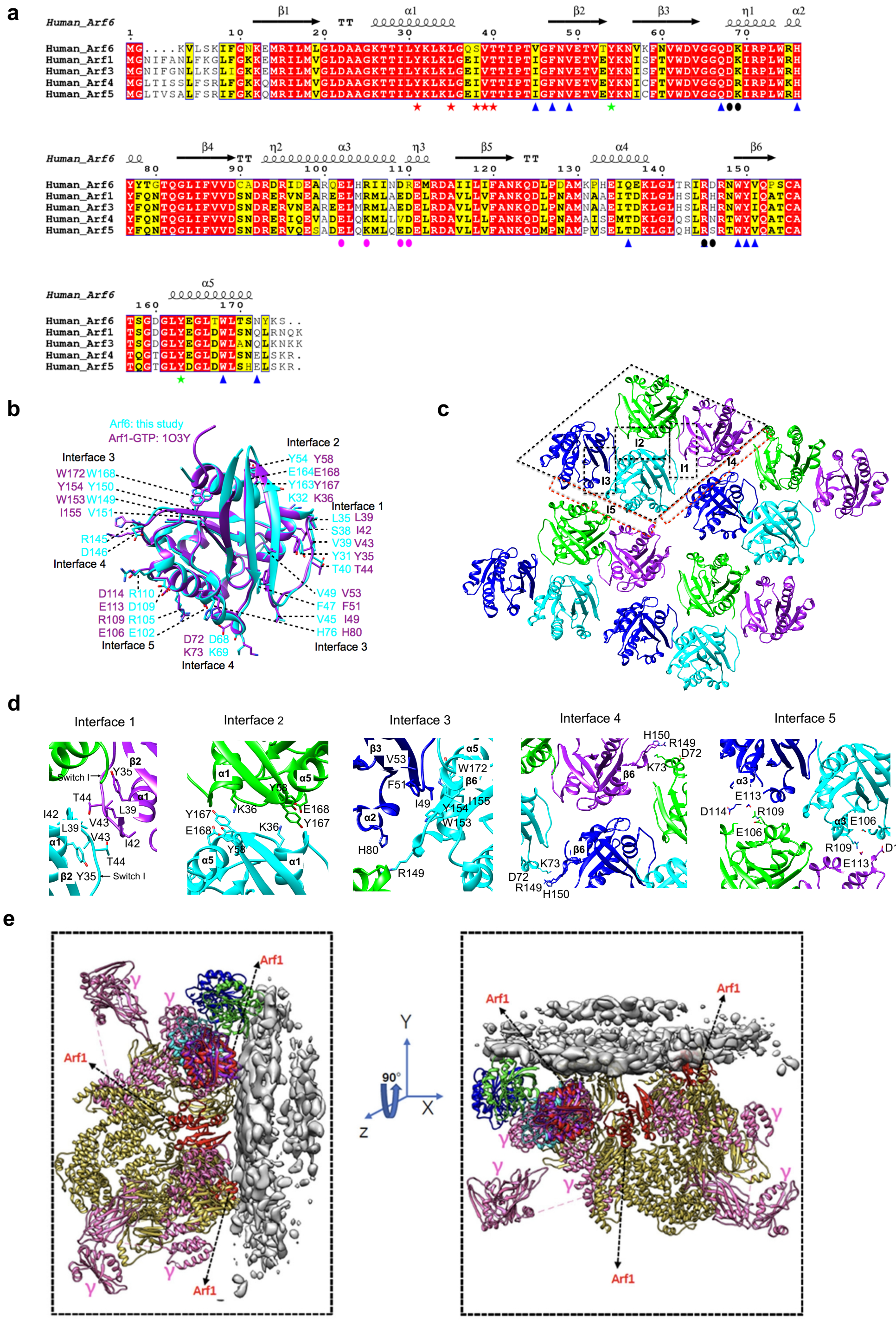

**Extended Data Fig. 4. Modelling of ARF1 onto the ARF6 lattice.**

**a**, Sequence alignment of human ARF6 with ARF1, ARF3, ARF4 and ARF5. The secondary structure assignments are derived from a model of ARF6 (PDB ID: 2J5X). Conserved and similar residues are shown in red and yellow, respectively.

Interfaces 1, 2 and 3 are labelled with red star, green star and blue triangle, respectively. Interfaces 4 and 5 are labelled with black and magenta circles, respectively. **b**, Structural comparison of ARF6-GTP on membrane with ARF1-GTP in solution (PDB ID: 1O3Y). The corresponding residues in ARF1 and ARF6 for the different interfaces of the lattice structures are indicated. **c**, Organization of ARF1 tetramers in helical arrays. The large black dashed box outlines four subunits forming a tetramer. The smaller black dashed boxes highlight the major protein interfaces. **d,** Major protein interfaces in the ARF1 lattice. Interface 1 is formed by helix α1 and switch I interacting with their counterpart in an adjacent subunit in an anti-parallel manner. Interface 2 is formed by helix α5 and inter-switch from adjacent subunits forming symmetric interaction. Interface 3 is formed by switch I, sheet β6 and helix α5 between two adjacent subunits forming interactions. Interface 4 involves tetramer interactions within the same helical row with predicted electrostatic interactions highlighted. Interface 5 involves tetramers on different helical rows interacting with each other with predicted salt bridges forming symmetric interactions highlighted. **e,** Comparison between the positioning of ARF1 in the lattice structure versus that in the ARF1-coatomer complex in the cryoEM structure of COPI vesicles (PDB code, 5NZR). The Arf1 molecules in the ARF1-coatomer complex are colored in pink, while they are colored in blue, green, cyan, and purple in the ARF1 tetramer of our lattice structure. The superimposition of one Arf1 molecule in the lattice structure (purple) on to one ARF1 in the ARF1-coatomer complex results in severe structural collision between another Arf1 molecule (cyan) of the lattice structure and the gamma-COP subunit (pink) of the ARF1-coatomer complex.

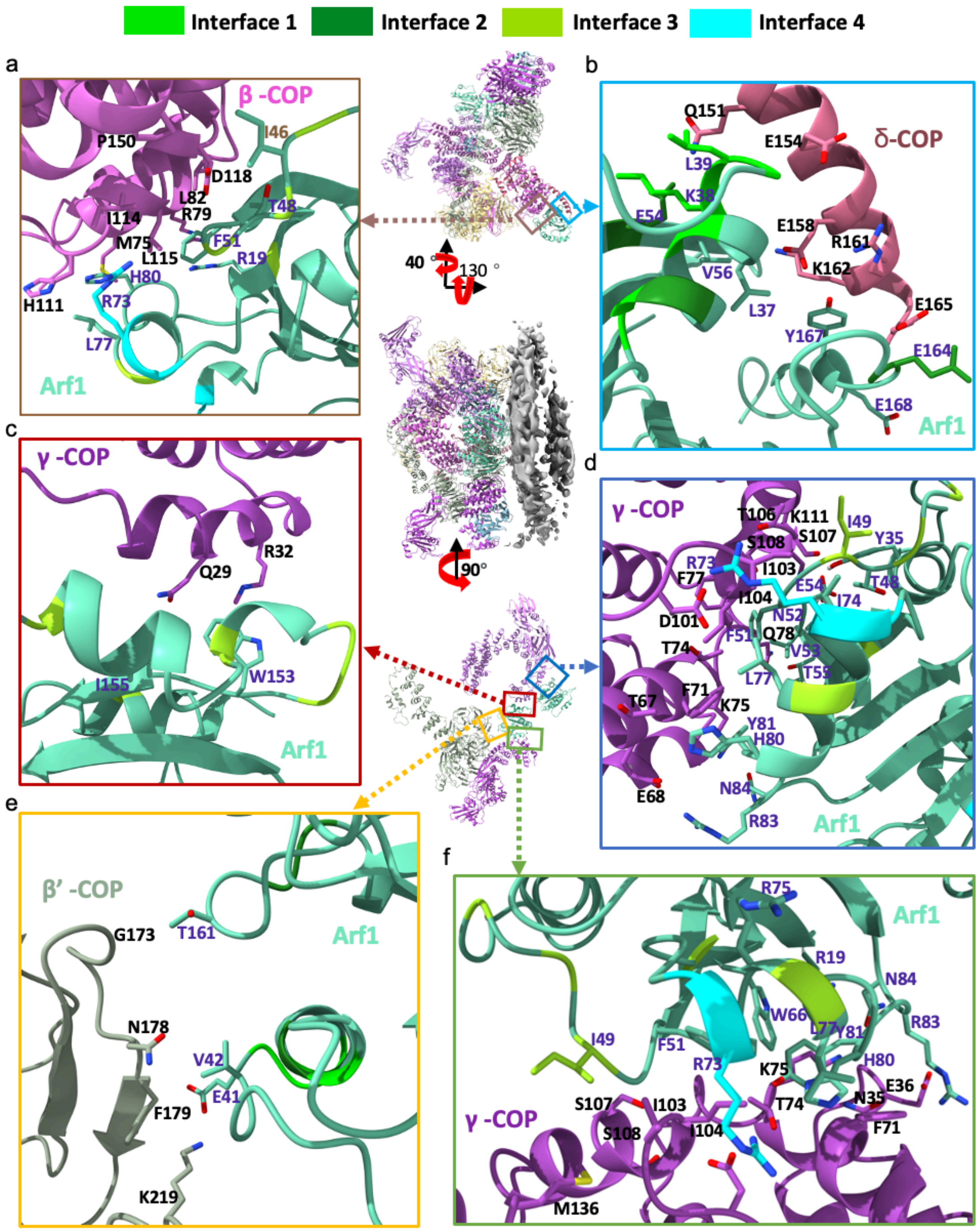

**Extended Data Fig. 5. Comparing the positioning of key residues in the protein interfaces of the ARF1 lattice to their positioning in the ARF1-coatomer complex on COPI vesicles.** Based on the 9-angstrom resolution cryoEM map as well as the reported molecular model (PDB code, 5NZR) of ARF1-

coatomer complex on the membrane, we built the full model with all residues assigned and then analyzed all the potential interfaces between ARF1 proteins (colored in Mint) and coatomer subunits, β-COP (a), δ-COP (b), β'-COP (e), and γ-COP (c,d,f). The potential residues of ARF1 in the interfaces are labeled in blue and that of coatomer subunits in black. Key residues involved in the protein interfaces of the ARF1 lattice are colored in chartreuse (interface 1), emerald (interface 2), pear (interface 3) and cyan (interface 4), respectively. As human ARF1 was used to build the ARF1 lattice and yeast (*Saccharomyces cerevisiae*) ARF1 was used to build the ARF1-coatomer complex, some residues of yeast ARF1 show conserved changes as compared to that of human ARF1.

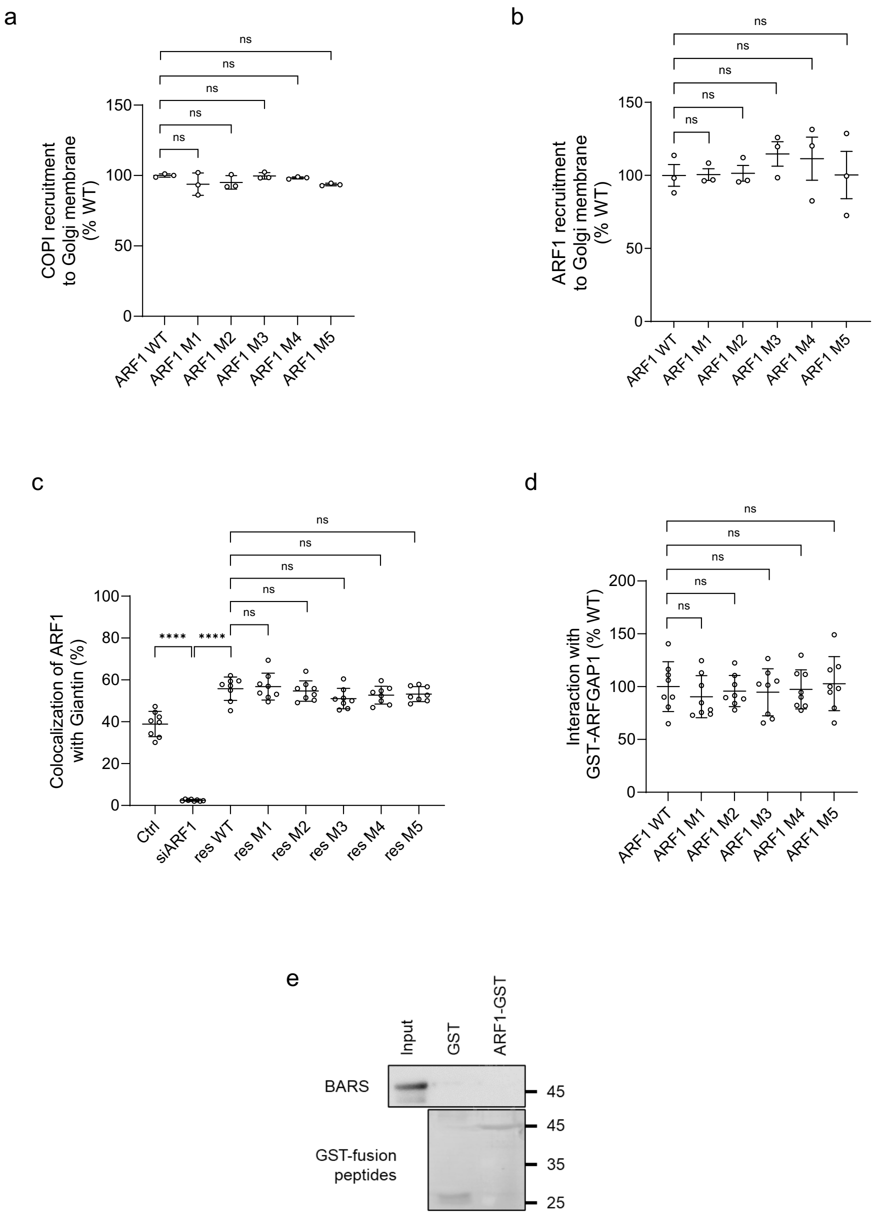

**Extended Data Fig. 6. Functional studies on the effect of ARF1 mutations.**

**a,** ARF1-mediated recruitment of coatomer to Golgi membrane assessed by reconstitution studies, n=3. Quantitation of three experiments is shown, ns ($p>0.05$), Student's t-test. **b,** The recruitment of ARF1 to Golgi membrane assessed by reconstitution studies, n=3. Quantitation of three experiments is shown, ns ($p>0.05$), Student's t-test. **c,** Golgi localization of ARF1 assessed by colocalization with a Golgi marker (giantin), n=3. Quantitation is shown for a representative experiment, ns (p>0.05), **** (p<0.0001) Student's t-test. **d**, Pulldown assay using purified components to assess ARF1 interacting with ARFGAP1, n=8. Quantitation of eight experiments is shown, ns (p>0.05), Student's t-test. **e,** Pulldown assay using purified components to assess ARF1 interacting with BARS, n=3. A representative result is shown.

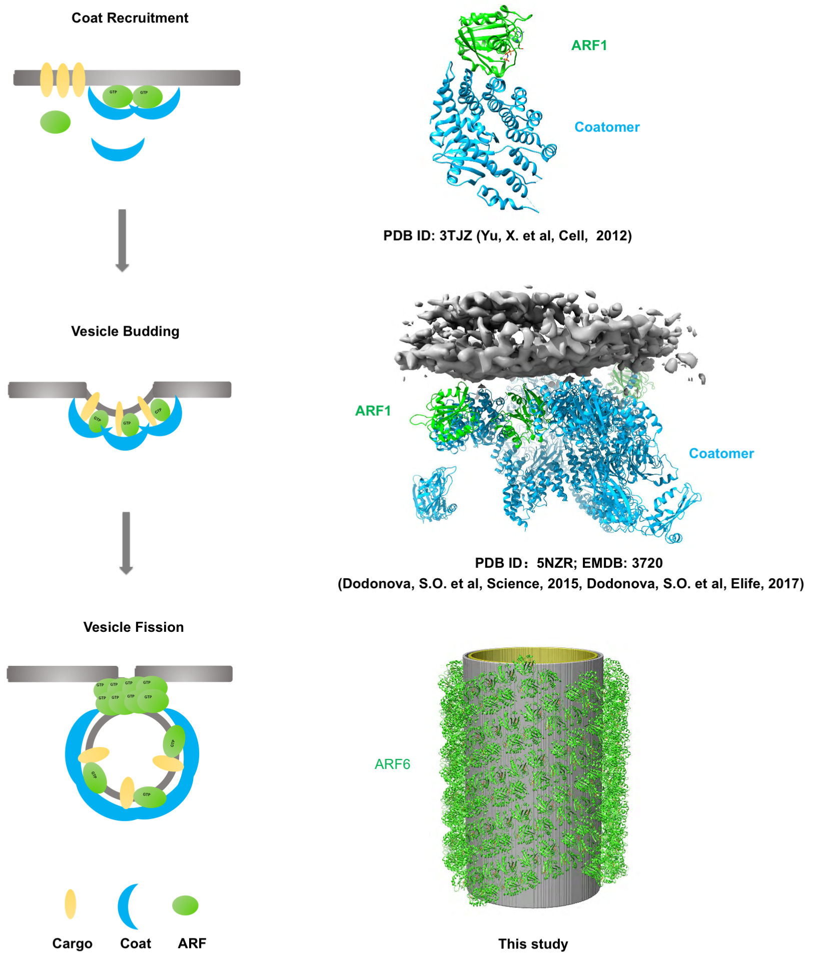

**Extended Data Fig. 7. Schematic of the major steps in vesicle formation.** The steps of coat recruitment, vesicle budding and vesicle fission are shown. Whereas molecular insight into how ARF acts in the first two stages have been achieved through structural studies, the current study provides molecular insight into the third stage through the structural elucidation of how ARF forms a tubular lattice structure on membrane.

**Extended Data Table 1:** Cryo-EM data collection, refinement, and validation statistics.

| | Arf6 in membrane binding state (EMD-33414) (PDB 7XRD) |
|---|---|
| **Data collection and processing** | |
| Magnification | 75,000 |
| Voltage (kV) | 300 |
| Electron exposure (e–/Å$^2$) | 50 |
| Defocus range (μm) | -1.5~-2.2 |
| Pixel size (Å) | 1.10 |
| Symmetry imposed | Yes |
| Initial particle images (no.) | 56557 |
| Final particle images (no.) | 20820 |
| Map resolution (Å) | 3.9 |
| FSC threshold | 0.143 |
| Map resolution range (Å) | 3.8-4.5 |
| **Refinement** | |
| Initial model used (PDB code) | 2J5X |
| Model resolution (Å) | 3.8/4.0 |
| FSC threshold | 0.38/0.5 |
| Model resolution range (Å) | 3.8 |
| Map sharpening *B* factor (Å$^2$) | -197 |
| Model composition | |
| Non-hydrogen atoms | 5464 |
| Protein residues | 656 |
| Ligands | 4 |
| *B* factors (Å$^2$) | |
| Protein | 49.28 |
| Ligand | 1.35 |
| R.m.s. deviations | |
| Bond lengths (Å) | 0.003 |
| Bond angles (°) | 0.976 |
| Validation | |
| MolProbity score | 0.9 |
| Clashscore | 1.56 |
| Poor rotamers (%) | 0.52 |
| Ramachandran plot | |
| Favored (%) | 97.99 |
| Allowed (%) | 2.01 |
| Disallowed (%) | 0 |

**Extended Data Table 2.** Statistics of the major protein interfaces in the ARF6 lattice assembly.

| Assembly | Interaction Interface | Interface area (Å$^2$) | ratio of interface area to total area |
|---|---|---|---|
| Inside tetramer | Interface 1 | 268 | 1/30 |
| | Interface 2 | 398 | 1/20 |
| | Interface 3 | 713 | 1/12 |
| Inter-tetramer, same row | Interface 4 | 86 | 1/96 |
| Inter-tetramer, different row | Interface 5 | 255 | 1/32 |

**Extended Data Table 3.** List of all mutations of ARF6 and ARF1 in this study.

| | Mutant | Site | Interface |
|---|---|---|---|
| ARF6 | M1 | Y31A/L35A/V39A | Interface 1 |
| | M2 | Y54A/Y163A | Interface 2 |
| | M3 | V45A/F47A/W149A/Y150A/V151A/W168A | Interface 3 |
| | M4 | K69E/R145E | Interface 4 |
| | M5 | R105E/R110E | Interface 5 |
| | M6 | F9W | N-terminal Amphipathic helix |
| | M7 | F9A | N-terminal Amphipathic helix |
| | M8 | F9E | N-terminal Amphipathic helix |
| | M9 | V4E/L5E/I8E | N-terminal Amphipathic helix |
| ARF1 | M1 | I42A/T44A | Interface 1 |
| | M2 | Y167A/E168R | Interface 2 |
| | M3 | I49A/F51A | Interface 3 |
| | M4 | K73E/R149E | Interface 4 |
| | M5 | R109E/D114R | Interface 5 |
| | M6 | F13E | N-terminal Amphipathic helix |

**Supplementary Video 1.** Overall ARF6 assembly on tubulated membrane.

**Supplementary Video 2.** Tetramer is the asymmetric unit of ARF6 helical packing.

**Supplementary Video 3.** Time frames of the molecular simulation for ARF6 tetramer interaction with lipid membrane.
